# Resilience-driven neural synchrony during naturalistic movie watching

**DOI:** 10.1101/2023.10.12.562025

**Authors:** Shuer Ye, Leona Rahel Bätz, Avneesh Jain, Alireza Salami, Maryam Ziaei

## Abstract

Psychological resilience protects individuals against the negative consequences of exposure to adversity. Despite increasing attention given to resilience for its role in maintaining mental health, a clear conceptualization of resilience remains elusive, and the intricacies of its neural correlates are poorly understood. Here, we recorded brain activity in healthy young adults using a 7T MRI scanner while they naturally watched movies. Stronger and more extensive resilience-driven neural synchrony, as estimated by inter-subject correlation, was observed in a wider set of brain regions in response to the negative movie compared to the neutral movie. Moreover, we found that high-resilience individuals had similar neural activities to their peers, while low-resilience individuals showed more variable neural activities. Intolerance of uncertainty (IU), a personality trait that shapes biased perception and cognition, damped the resilience-driven brain synchrony in regions related to attention, indicating IU may compromise resilience by affecting attentional functions. We propose that similarity of neural responses among resilient individuals highlights adaptive emotional processing. Conversely, the variability in neural responses indicates vulnerability to adverse psychological outcomes. These insights shed light on the mechanisms of resilience, highlighting that it operates as a system encompassing multiple neuropsychological processes crucial for adapting to external stimuli.

## Introduction

Psychological resilience, the ability to cope effectively with adversities, plays a crucial role in determining individuals’ overall well-being(Feldman, 2020), especially in the contemporary world where traumatic and highly stressful events frequently occur(Gershon et al., 2013; Pfefferbaum & North, 2020). Although conceptualized as a stable trait for decades, researchers increasingly recognize resilience as a dynamic process encompassing positive adaptation in the face of negative events(Denckla et al., 2020; Rutter, 2012; Vella & Pai, 2019). Behavioral studies have demonstrated that individual differences in resilience are closely linked to how individuals perceive, recognize, and process external stimuli(Parsons et al., 2016; Schäfer et al., 2015). Individuals with higher resilience tend to report more positive emotional experiences(Tugade et al., 2004), demonstrate quicker disengagement from emotional stimuli(Yi et al., 2020), and employ adaptive emotional regulation strategies more effectively(Polizzi & Lynn, 2021). Additionally, imaging studies provide evidence that individual differences in resilience may stem from functional variations in brain networks involved in top-down attentional processing(Jääskeläinen et al., 2021), salience detection(Homberg & Jagiellowicz, 2022), socio-emotional functioning(Liu et al., 2023), and cognitive control(Rodman et al., 2019). These networks are pivotal in sustaining individuals’ resilience and in facilitating adaptive behavior after exposure to stressors(Brunetti et al., 2017; Roeckner et al., 2021). From a neuropsychological perspective, resilience can be characterized as a dynamic and complex system that spans multiple psychological processes rooted in distinct underlying neural pathways. However, there remains a significant gap in neurobiological evidence to substantiate this characterization.

Such a complex system can be described by the Anna Karenina (AnnaK) principle(Moore, 2001; Zaneveld et al., 2017a), which highlights the vulnerability of a large, sophisticated system in a way that the failure of a single component can lead to the collapse of the entire system. Under this framework, the malfunctioning of one or more components within the resilience system leads to reduced resilience, making individuals susceptible to affective disorders. Essentially, individuals exhibit robust and effective resilience only when they consistently demonstrate adaptability throughout their interactions with negative events. Accordingly, those with high resilience typically show similar patterns of behavior and brain activity. In contrast, individuals with lower resilience display varied responses due to varying deficits in emotional and cognitive functions when faced with adversity. However, no study has investigated the neural mechanisms of resilience involved when participants are exposed to naturalistic and dynamic stimuli in experimental settings.

In this study, we aim to provide evidence for how the resilience process is reflected in brain activity during real-life experiences by using movie-fMRI(Finn, 2021; Meer et al., 2020; Sonkusare et al., 2019). By recording brain activity while participants watch movies, inter-subject correlation (ISC) can be calculated to measure the similarity of brain activity in response to the same external stimuli. Higher ISC indicates greater neural synchrony, suggesting that individuals process information in a similar manner(Hasson et al., 2010). Previous research on psychiatric population involving movie-fMRI has shown a decrease in the similarity of neural responses among patients with attention deficits and hyperactivity disorder(Salmi et al., 2020), schizophrenia(Tu et al., 2019), and depression(Gruskin et al., 2020). Recent investigations also suggest socially well-connected individuals, who typically show higher levels of psychological well-being(Nummenmaa et al., 2018; J. Wang et al., 2018), tend to exhibit similar brain activity patterns while watching movies(Baek et al., 2022; Guthrie et al., 2022). Building upon these findings, it is reasonable to infer that convergent or similar neural activities during movie-watching may serve as indicators of good mental states and psychological well-being (i.e., high resilience).

We further explored how intolerance of uncertainty (IU), a dispositional trait indicating a tendency to avoid uncertainty and harbor negative beliefs about ambiguous or uncertain situations(Carleton, 2012; Grenier et al., 2005), could impact the resilience system(Tanovic et al., 2018). IU plays a vital role in gating attentional engagement(Dieterich et al., 2016; Sahib et al., 2023) and shaping emotion perception(Perlman, 2009), and it is a strong risk factor for generalized anxiety disorder(Gu et al., 2020). Therefore, IU would primarily undermine the resilience system via biased emotion perception and abnormal attention selection, essential processes that support the resilience system.

Our primary objectives are twofold: first, to scrutinize the impact of resilience on individuals’ neural responses by employing naturalistic stimuli and intersubject representational similarity analysis (IS-RSA)(G. Chen et al., 2017), an innovative ISC-based approach to robustly detect individual differences and the underlying brain dynamics in naturalistic settings. We hypothesize that individuals with high resilience display more consistent neural responses, while those with low resilience exhibit more idiosyncratic neural responses, according to the AnnaK model. Second, to identify the modulatory role of IU in the relationship between resilience and neural synchrony. We anticipate that IU will attenuate the neural synchrony that is driven by resilience, especially in the brain areas that are responsible for attention and perception.

## Materials and Methods

### Participants

Seventy-eight younger adults were recruited from the local community via paper flyers and online advertisements to participate in this study. Due to technical issues, incomplete data, and excessive movement (i.e., average framewise displacement > 0.25 mm), 62 participants (33 males and 29 females, age range 19 to 35 years, mean age 25.68 ±4.30 years) were included for further data analyses. All participants provided written informed consent and were compensated with a gift card for their participation. All participants were healthy, right-handed, and reported no history of a neurological (e.g., epilepsy, stroke, or brain injury) or psychiatric diseases (e.g., depression and anxiety, schizophrenia, or autism spectrum disorder) over the past five years. They were not taking any medications for mood disorders at the time of experiment. The study was approved by the regional ethics committee (approval #390390).

### Procedure

The study consisted of two sessions: a brain imaging session and a behavioral session (**Fig. 1A**). First, participants took part in an MRI session, which included both functional and structural brain imaging protocols. Following the brain imaging sessions, the behavioral session took place within two days and lasted about two hours, involving self-reported questionnaires and computerized tasks.

**Figure 1.**
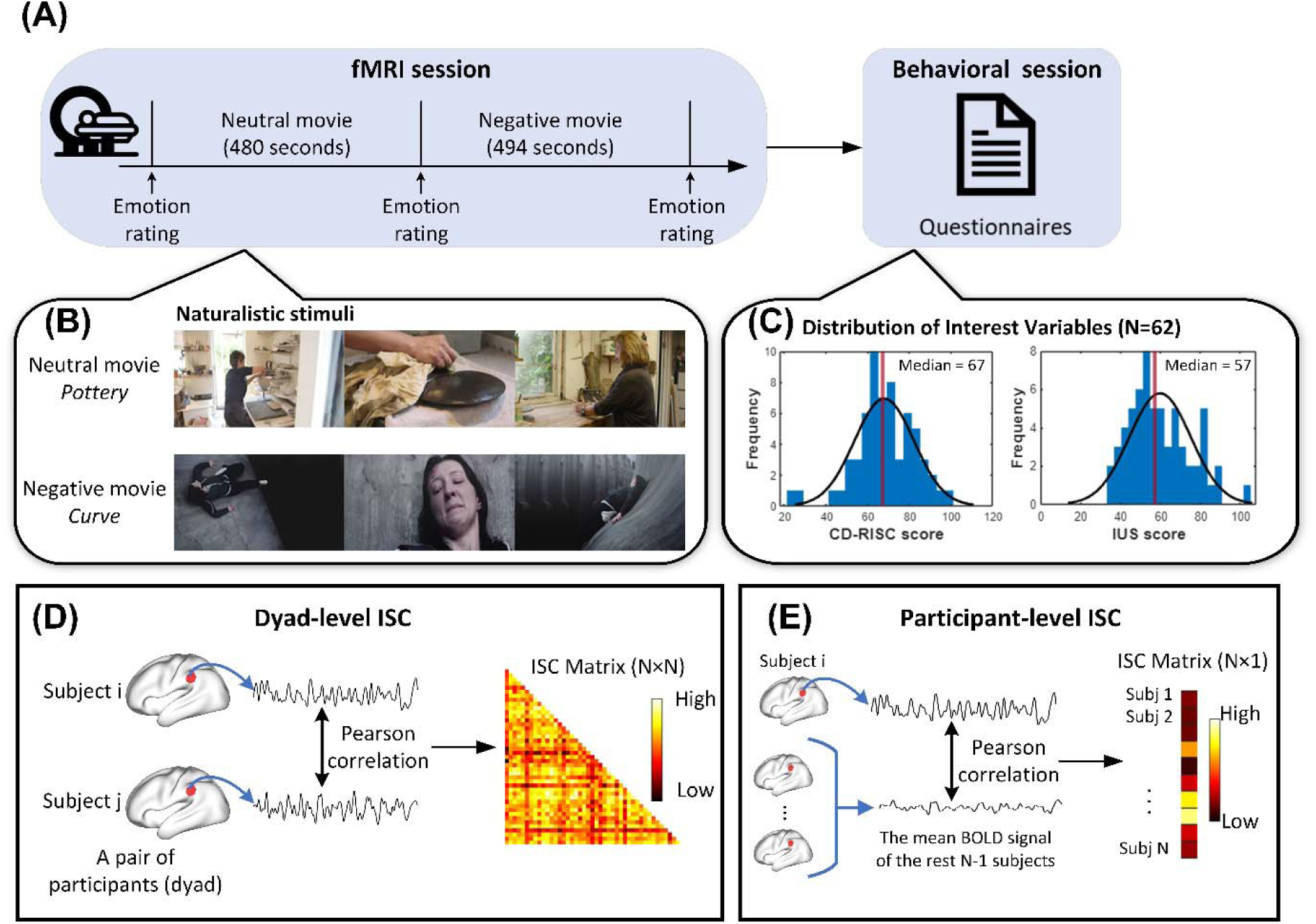
Study paradigm and calculation of inter-subject correlation (ISC). (A) Participants first underwent the fMRI scanning while they watched two movies. Before and after each movie, they reported their emotional valence and arousal. After the fMRI session, participants were invited back to complete a behavioral session comprising of self-reported questionnaires and cognitive testing as part of the protocol for a larger study on resilience and aging brain. (B) Participants first watched the neutral movie, “*Pottery*”, and then the negative movie, “*Curve*”, in the 7T MRI scanner. (C) Histogram plots for resilience score measured by Connor-Davidson Resilience Scale (CD-RISC) and intolerance to uncertainty score measured by Intolerance of Uncertainty Scale (IUS). Scores of both questionnaires are normally distributed. (D) Dyad level ISC was computed by correlating the BOLD signal time series of the two participants within each dyad. (E) Participant-level ISC was computed by correlating the BOLD signal time series of the given participant and the mean BOLD signal time series of the rest of participants. The inferior parietal lobe is depicted here as an example for the analyses.

Before the brain imaging session, participants received instructions about the procedure and the experiment, both verbally and in a written format. During the scan, participants underwent an 8-minutes anatomical scan followed by a movie watching task. They were presented with a neutral movie, followed by a negative movie, and were asked to rate their emotional valence and arousal levels before and after each movie clip by verbally reporting their response on a 9-point Self-Assessment Manikin scale. Employing fixed movie sequences ensured consistent stimulus context across participants and effectively eliminates confounding factors arising from variability in stimulus presentation(Baek et al., 2022; P.-H. A. Chen et al., 2020). Participants were instructed to watch the videos naturally while trying to remain still during the scanning session. Foam padding was used to minimize head motion, and participants listened to the videos’ sound through earphones (BOLDfonic, Cambridge Research Systems Ltd) compatible with 7T MRI scanner while watching the videos through a mounted mirror on the head coil.

### Movie clips

Prior to the main study, a pilot behavioral study was conducted to select materials for the main experiment. Four movies from prior studies and online repository(Taylor et al., 2017) were chosen for the pilot experiment. Twenty-four younger adults provided a series of ratings about valence, arousal, and emotion types, as well as a continuous intensity rating, for each movie clip. Based on the results of the pilot study, two movie clips were chosen for the subsequent brain imaging study (more details of the pilot study can be found in **Supplementary information**, **Fig.S1**). The neutral movie, titled “*Pottery*”, depicts two women engaged in pottery-making, which lasted for 480 seconds. The negative movie, named “*Curve*”, tells the story of a woman struggling to maintain her balance on the slope of a dam, desperately trying to avoid falling into the abyss below, which lasted for 494 seconds (**Fig. 1B**).

### Questionnaires

As this study was part of a larger study, participants were asked to fill out a variety of self-reported questionnaires including anxiety, depression, and perceived stress, and to complete computerized tasks including theory of mind, stroop, and emotion perception.

The level of psychological resilience was assessed by using the Connor-Davidson Resilience Scale (CD-RISC), which is a well-validated scale to screen individuals for varying level of resilience(Connor & Davidson, 2003) and has been used in variety of settings including behavioral(van Gils et al., 2022), clinical(Hendricks et al., 2023), and neuroimaging studies(Long et al., 2019). The scale consists of 25 items such as ‘I am able to adapt when changes occur’. Each item is scored on a 5-point Likert scale from 0 (Not true) at all to 4 (True nearly all the time). Higher scores indicate a higher level of resilience. In the current study, the Cronbach’s Alpha of the CD-RISC is 0.902, reflecting high internal consistency reliability in our sample.

The intolerance of uncertainty was assessed with the Intolerance of Uncertainty Scale (IUS)(Freeston et al., 1994), which comprised of 27 items such as ‘Unforeseen events upset me greatly’, on a 5-point Likert scale from 1(Not at all characteristic of me) to (5-Entirely characteristic of me). Higher scores indicate increased intolerance of uncertainty. The Cronbach’s Alpha of the IUS is 0.904 in the current sample. Furthermore, we utilized the Hospital Anxiety and Depression Scale(Bjelland et al., 2002), Perceived Stress Scale(Lee, 2012), and Difficulties in Emotion Regulation Scale (DERS) (Gratz & Roemer, 2004) to evaluate participants’ levels of anxiety, depression, and stress, as well as their difficulties in emotion regulation.

### Imaging data acquisition

The imaging data were collected on a 7T MRI Siemens MAGNETOM Terra scanner with a 32-channel head coil. Functional images were acquired using a multi-band accelerated echo-planar imaging sequence (92 interleaved slices, multi-band acceleration factor□=□8, echo time = 19 ms, repetition time = 2000 ms, matrix size = 160 × 160 mm, voxel size = 1.25 × 1.25 ×1.25 mm, field of view = 200 mm, slice thickness = 1.25 mm, and flip angle = 80°), resulting in 243 volumes for the neutral movie and 250 volumes for the negative movie.

The structural T1-weighted high-resolution scans were acquired using a MP2RAGE sequence (224 sagittal slices, echo time = 1.99 ms, repetition time = 4300 ms, inversion time 1 = 840 ms, inversion time 2 = 2370 ms, voxel size = 0.8 × 0.8 × 0.8 mm, flip angle = 5°/6° and slice thickness = 0.75 mm).

### Imaging data preprocessing

The imaging data underwent preprocessing using fMRIPrep version 22.0.2(Esteban et al., 2019), followed by post-preprocessing using XCP-D version 0.3.0(Ciric et al., 2018). The preprocessing pipeline included slice-time correction, head-motion correction, co-registration between structural and functional images, normalization to MNI space, nuisance regression (i.e., the top 5 principal aCompCor components from white matter and cerebrospinal fluid compartments and the six motion parameters and their temporal derivatives), band-pass filtering within the 0.009-0.08 Hz, and smoothing of the images using 4mm Kernel size. As part of the preprocessing, the first three dummy scans were discarded, leaving 240 volumes for the neutral movie and 247 volumes for negative movie conditions. Furthermore, Region of interest (ROI) signal extraction was conducted based on the Schaefer 200-parcel cortical parcellation(Schaefer et al., 2018) and Tian 16-parcel subcortical parcellation(Tian et al., 2020), resulting in time series from 216 brain regions for each condition and for every participant. These 216 regions were assigned to eight spatial independent brain networks (i.e., the visual network [VN], somatomotor network [SMN], dorsal attention network [DAN], ventral attention network [VAN], control network (CN), default mode network [DMN], Limbic network [LN], and subcortical network [SUB]).

### Inter-subject correlations

To test whether both neutral and negative movies elicited robust neural synchrony across all participants, we calculated the participant-level intersubject correlation (ISC) by correlating the BOLD signal time series of each participant with the mean BOLD signal time series of the rest of participants (**Fig. 1E**). This procedure was conducted for each ROI and each participant, resulting in a 216 ROI × 62 participant-level ISC matrix. Furthermore, Pearson’s correlation and paired t-test were performed to compare the ISC between neutral and negative conditions, allowing us to estimate the similarity and variation of brain synchrony between the two conditions.

The dyad-level ISC was calculated for each dyad (pair of participants) to indicate pairwise brain synchrony during movie-watching (**Fig. 1D**). To construct the ISC matrix, we computed Pearson’s correlation between the BOLD signal time series in each of the 216 brain regions for each pair of participants. With a total of 1891 unique dyads generated from the 62 participants included in the study, this procedure resulted in two 216×1891 ISC matrices separate for neutral and negative movies. Subsequently, the ISC matrices were transformed using Fisher’s *r* to *z* transformation to increase data normality.

### Intersubject representational similarity analyses

In the current study, we conducted three IS-RSA using linear mixed-effects (LME) models with crossed random effects(G. Chen et al., 2017; Finn et al., 2020) to examine our hypotheses. To build these models, we utilized LME4 (version 1.1-32) and LMERTEST (version 3.1.3) in R(Baek et al., 2022). The LME models were advantageous as they could account for the random effects and correlation structure of unique pairs of participants within dyads, effectively addressing the statistical dependencies between pairwise observations(Bernal-Rusiel et al., 2013; G. Chen et al., 2017).

To test the first hypothesis that resilience-driven brain synchrony may follow the AnnaK model, with high-resilience individuals exhibiting similar neural responses while low-resilience individuals having idiosyncratic neural responses, we performed IS-RSA with binarized resilience scores. We first divided all participants into high resilience and low resilience groups based on the median of the resilience score (**Fig. 3A**). This characterization resulted in three types of dyads among the 1891 unique pairs: (1) {high, high} for dyads where both participants were in the high resilience group; (2) {high, low} for dyads with one high resilience and one low resilience participant; (3) {low, low} for dyads with both participants in the low resilience group. Subsequently, we conducted the first IS-RSA by constructing LME models, incorporating random effects of participants within each dyad. The model included the binarized dyad-level resilience as a fixed effect and the dyad-level ISC for a given brain region as the dependent variable. A planned-contrast analysis was then used to examine and compare the neural similarity among the three types of dyads.

To explore the first hypothesis in a greater detail, we further performed the second IS-RSA with resilience similarity as the independent variable. Based on the AnnaK model(Ohad & Yeshurun, 2023), The resilience similarity was defined by the minimum resilience score for each dyad. Thus, only pairs of high-resilience individuals show high similarity (i.e., high-resilience individuals are alike), whereas pairs of low-resilience or mixed pairs (a pair consists of one with low resilience and one with high resilience) were consistently characterized by low similarity (i.e., low resilience individuals differ from their peers as well as high-resilience individuals) (**Fig. 3C**). To ensure the normality of the data, log-transform was further applied to resilience similarity scores. This allowed us to test whether resilience similarity could predict the neural similarity observed in dyads. By doing so, we aimed to gain deeper insights into how the similarity in resilience scores of dyads relates to the extent of neural similarity observed in their brain responses.

Lastly, we further examine our second hypothesis and investigate whether IU affects resilience-driven neural synchrony. The product of the IU score within each dyad was normalized to a 0-1 range, resulting in joint intolerance of uncertainty score (JIUS), indicating both the similarity of IU between two individuals within the dyads and the absolute position of the dyad members on the IU spectrum (i.e., the level of IU). We performed the third IS-RSA, which investigated the modulating effect of IU by introducing the interaction effect between resilience similarity and JIUS.

For each IS-RSA, false discovery rate (FDR) correction with *p* < 0.05 was applied to control for multiple comparisons.

### Validation analysis

Given the similarity in neural responses is associated with demographic factors(Finn et al., 2020; Nummenmaa et al., 2018), we validated our results and accounted for the potential impact of demographic similarity on brain synchrony. In this analysis, we added the similarity of age, gender, and education as covariates into the LME models to assess the robustness of our findings. For gender, we employed an indicator variable to represent the gender similarity of each dyad, with 1 indicating the same gender and 0 indicating different genders. Regarding age, we calculated the L1 distance between the ages of each pair of participants. The L1 distance was then normalized into a range between 0 to 1, and the resulting value was subtracted from 1 to obtain a measure indicating the similarity in age. The similar procedure was conducted to obtain the similarity in education.

## Results

### Movies effectively trigger corresponding emotion experiences

Participants reported their emotional valence and arousal levels before and after each movie clip. After watching the neutral movie, no significant changes were observed in emotional valence (*t* = 0.095, *df* = 61, *p* = 0.925, *d* = 0.012) and arousal (*t* = 0.375, *df* = 61, *p* = 0.709, *d* = 0.048) compared with the baseline, prior to watching the movie. However, after watching the negative movie, compared to the neutral clip, a significant decrease in valence (*t* = –9.193, *df* = 61, *p* < 0.001, *d* = 1.167) and an increase in arousal (*t* = 7.298, *df* = 61, *p* < 0.001, *d* = 0.927) were observed (**Fig. 2**). These results demonstrate the validity of our naturalistic stimuli. Furthermore, no significant correlations were found between resilience score and emotion ratings (all P’s>0.075) nor resilience score and alteration of emotional status (all P’s > 0.1)).

**Figure 2.**
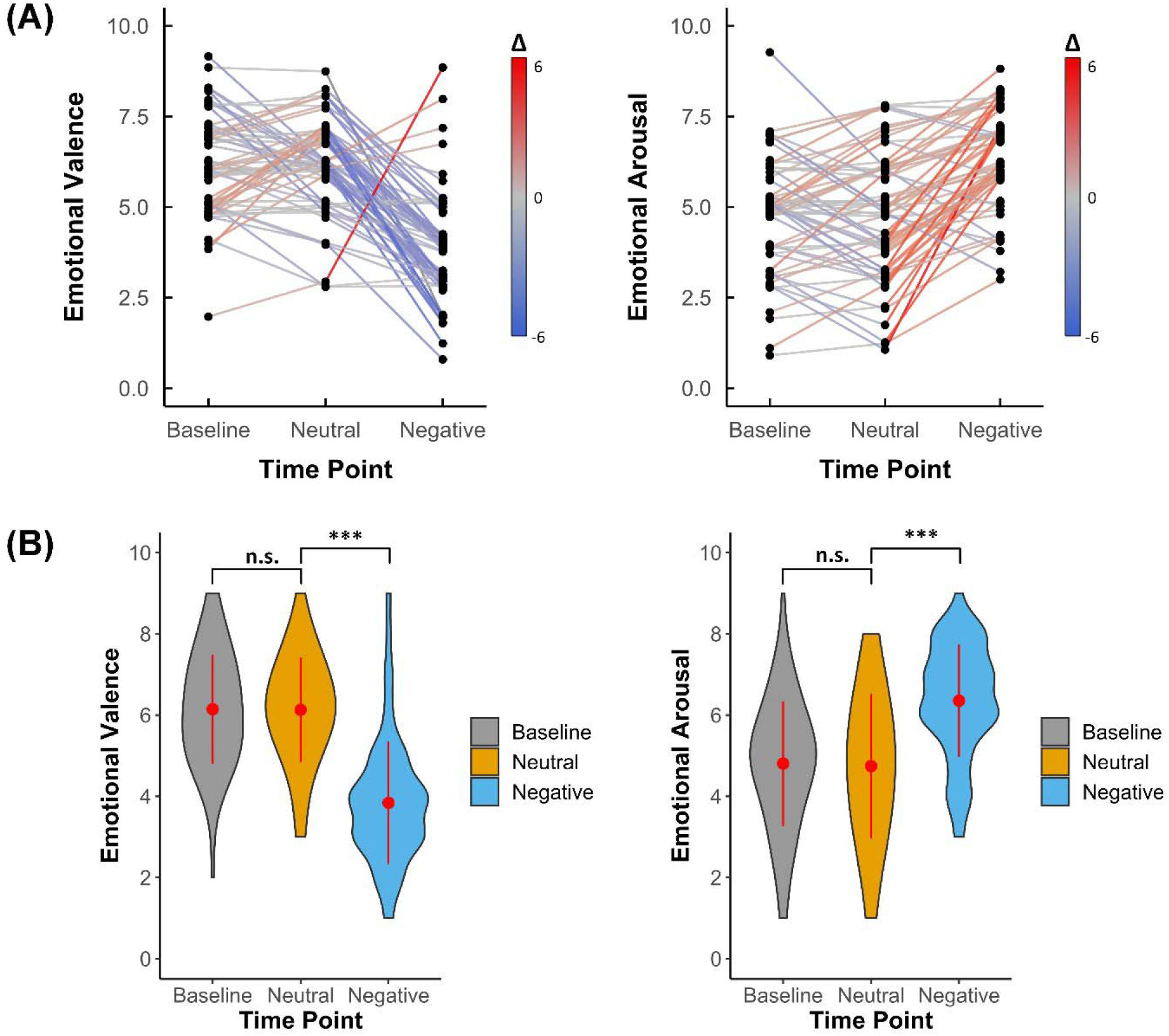
Emotional valence and arousal ratings before and after viewing the neutral and negative movies (participants N=62). (A) The emotional valence and arousal ratings for the before (baseline) and after each movie. The colors of the lines indicate changes in individual responses for the emotional ratings. (B) The violin plots of the emotional valence and arousal ratings. No significant differences were observed between baseline and after viewing the neutral movie in emotional valence, and the same results were obtained for emotional arousal. However, a significant decrease in emotional valence and a significant increase in emotional arousal were observed after participants watched the negative movie. The red points within the violin plots indicate the mean value of observation, and the red lines indicate the standard deviations. n.s.: not significant; ***: *p*<0.001.

### Low resilience is associated with low emotional well-being

We investigated the relationship of resilience with various mental health indicators. Negative correlation between resilience and IU (*r* = –0.395, *p* = 0.001), anxiety (*r* = –0.314, *p* = 0.013), depression (*r* = –0.306, *p* = 0.016), and perceived stress (*r* = –0.489, *p* < 0.001) were found. Moreover, negative associations were found among resilience and multiple subscales in DERS, including nonacceptance of emotional response (*r* = –0.401, *p* = 0.001), lack of emotional awareness (*r* = –0.423, *p* < 0.001), limited access to emotion regulation strategies (*r* = –0.377, *p* = 0.002), and lack of emotional clarity (*r* = –0.404, *p* = 0.001). Full list of zero-order correlations of resilience, IU, and related variables, including depression, anxiety, perceived stress, and emotion regulation difficulties are presented in the **Tab.1**. In summary, our behavioral data suggests that low resilience individuals may show a higher risk of mood disorders signifying their vulnerability to different emotional and cognitive dysfunctions.

**Table 1.** Pearson’s correlations between interested variables (N = 62). The correlation analysis was performed to illustrate the relationship between resilience, intolerance of uncertainty (IU), anxiety, depression, and difficulties in emotion regulation. The resilience was measured by the Connor-Davidson Resilience Scale. IU was assessed by the Intolerance of Uncertainty Scale. Anxiety and depression were measured by the Hospital Anxiety and Depression Scale. Stress was measured with the Perceived Stress Scale. Emotion regulation problems were evaluated by the Difficulties in Emotion Regulation Scale (DERS). IU: intolerance of uncertainty; DERS: Difficulties in Emotion Regulation Scale; The DERS subscales from 1 to 6 represent the following difficulties in emotion regulation: 1-Nonacceptance of emotional responses; 2 – Difficulty engaging in goal-directed behavior; 3-Impulse control difficulties; 4 – Lack of emotional awareness; 5 – Limited access to emotion regulation strategies; 6 – Lack of emotional clarity. *: p < 0.05; **: p < 0.01.

|  | 1 | 2 | 3 | 4 | 5 | 6 | 7 | 8 | 9 | 10 |
| --- | --- | --- | --- | --- | --- | --- | --- | --- | --- | --- |
| 1. Resilience | 1 |  |  |  |  |  |  |  |  |  |
| 2. IU | -.395** | 1 |  |  |  |  |  |  |  |  |
| 3. Anxiety | -.314* | .537** | 1 |  |  |  |  |  |  |  |
| 4. Depression | -.306* | .373** | .449** | 1 |  |  |  |  |  |  |
| 5. Stress | -.489** | .628** | .588** | .531** | 1 |  |  |  |  |  |
| 6. DERS |  |  |  |  |  |  |  |  |  |  |
| DERS-1 | -.401** | .287* | .421** | .197 | .422** | 1 |  |  |  |  |
| DERS-2 | -.248 | .275* | .220 | .396** | .384** | .180 | 1 |  |  |  |
| DERS-3 | -.162 | .382** | .191 | .315* | .403** | .178 | .319** | 1 |  |  |
| DERS-4 | -.423** | .112 | .060 | .313* | .276* | .149 | .017 | .246 | 1 |  |
| DERS-5 | -.377** | .353** | .278* | .434** | .543** | .507** | .414** | .460** | .359** | 1 |
| DERS-6 | -.404** | .300* | .180 | .390** | .436** | .300** | .172 | -.018 | .599** | .457** |

### Negative movie elicits stronger neural synchrony

To assess the robustness of neural synchrony elicited by both natural and negative movies across all participants, we computed the participant-level ISC for each brain region.

The results indicated robust overall brain synchrony for both movies, with the most pronounced effects in auditory and visual brain areas (**Fig.3A**). Higher ISC values were evident in the temporal lobe, prefrontal cortex, posterior cingulate cortex, precuneus, and subcortical areas (such as the hippocampus, amygdala, and caudate) during the negative movie as compared to the neutral movie (**Fig.3B**, **Tab.S2**). Overall, participants exhibited higher levels of brain synchrony during the negative condition compared to the neutral condition (mean ISC of neutral movie = 0.191, mean ISC of negative movie = 0.291, *t* = 23.255, *df* = 217, *p* < 0.001, *d* = 1.582, **Fig.3C**). Additionally, the spatial distribution of neural synchrony was highly correlated across the two conditions (*r* = 0.832, *p* < 0.001, **Fig.3D**), suggesting a stable functional organization in response to naturalistic stimuli regardless of the emotional content of the movies. In summary, the negative movie induced stronger neural synchrony compared to the neutral movie, especially in the areas that implicated in affective function (e.g., the insula and amygdala), social cognition (e.g., the prefrontal cortex and inferior parietal lobe) and memory retrieval (e.g., the posterior cingulate cortex and hippocampus).

**Figure 3.**
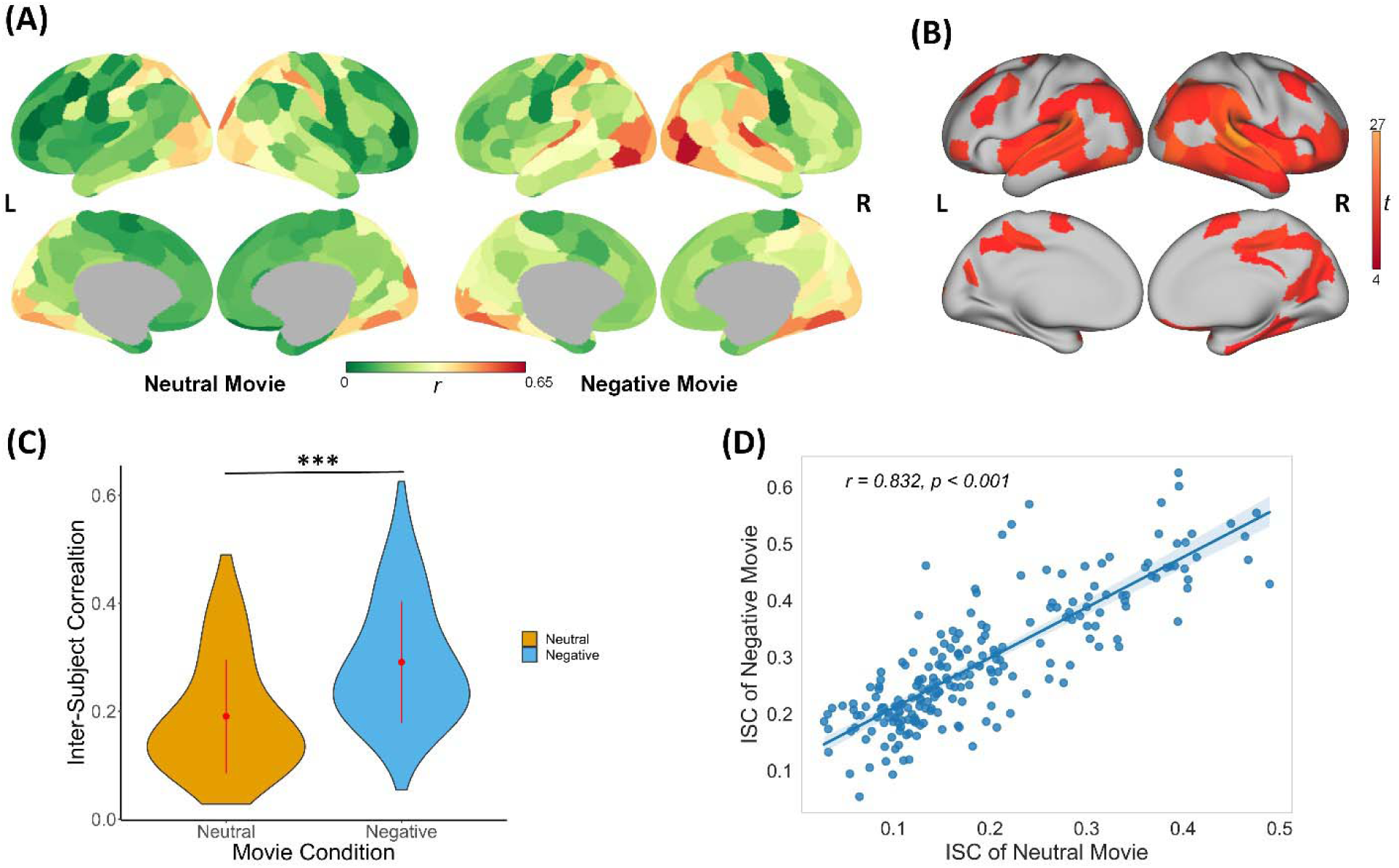
Participant-level inter-subject correlation (ISC) of the neutral and negative movies. (A) **t**he participant-level ISC maps for neutral and negative movies. (B) results of the paired t-test (FDR corrected) comparison between ISC of neutral and negative movie. The higher *t*-values indicates higher ISC elicited during the negative compared to the neutral movie. (C) the negative movie induced overall greater brain synchrony than the neutral movie. The red points within the violin plots indicate the mean value of the observation, and the red lines indicated the standard deviations. (D) the spatial distribution of ISC maps of two conditions is highly consistent. Each data point represents one brain area. The shaded zone represents 95% Confidence Interval (CI). ***: *p*<0.001.

### Resilience drives neural synchrony in a convergent way

To test the first hypothesis regarding whether individuals with higher resilience scores exhibit higher neural similarity during movie watching, we conducted a comparison between high and low resilience individuals using the IS-RSA with dyad-level ISC.

Participants were categorized into two groups based on their resilience scores, resulting in three types of dyads based on the resilience group membership of the pairs: {high, high}, {high, low}, and {low, low}. During the neutral movie, higher ISCs were found in {high, high} dyads compared to the {low, low} dyads in some regions, encompassing the postcentral gyrus, frontal eye fields, and ventral prefrontal cortex **(Fig.4B, Tab.S3)**. We also found higher ISCs in the {high, high} dyads than {high, low} dyads in the postcentral gyrus, lateral prefrontal cortex, and frontal eye fields. No significant results were found in the comparison of {high, low} and {low, low} dyads.

**Figure 4.**
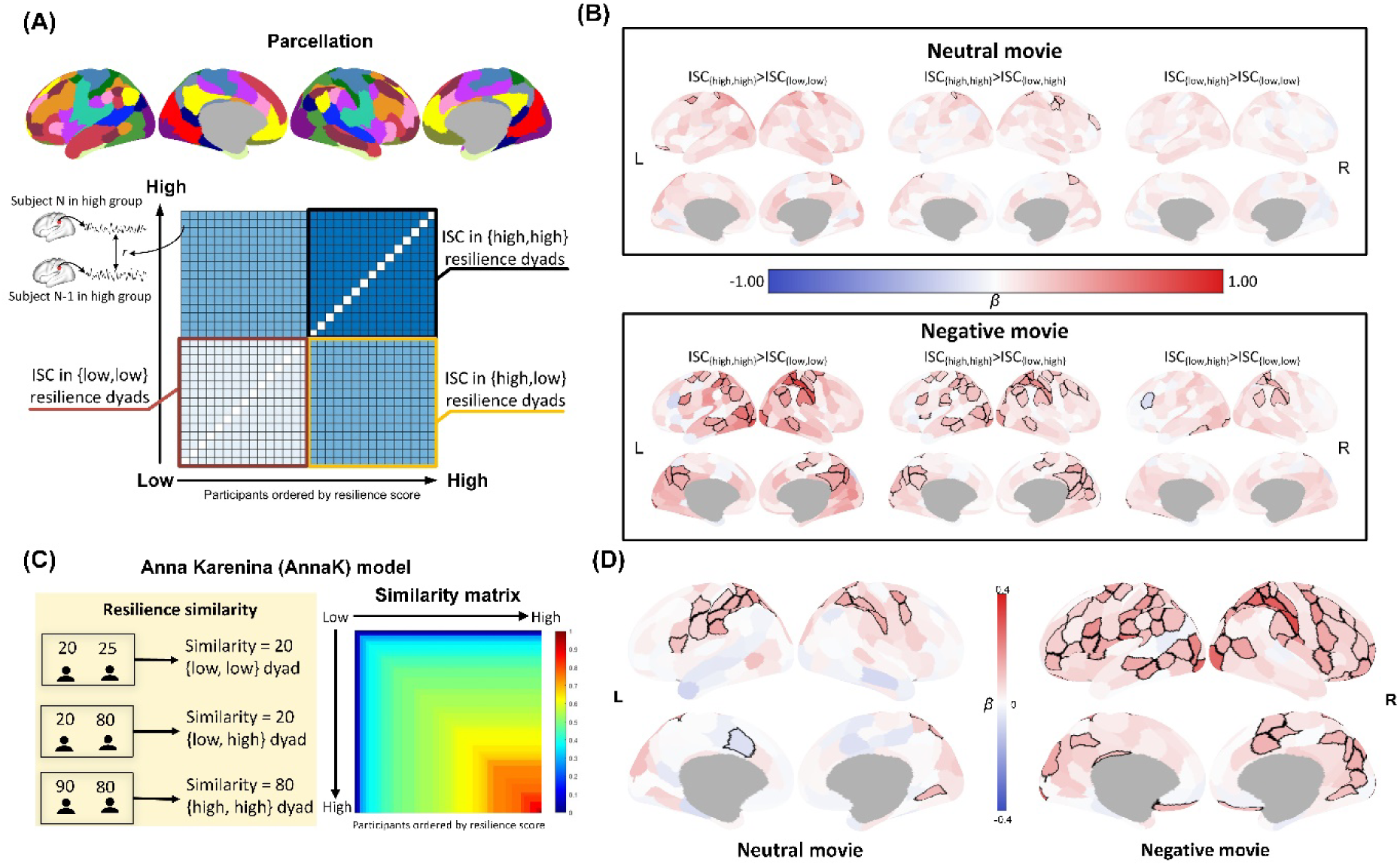
Results of dyad-level analyses on the relationship between resilience and neural synchrony. (A) All dyads were classified into three types, {high, high}, {high, low}, and {low, low}, based on the resilience levels of their respective members. The dyad-level Inter-Subject Correlation (ISC) was then calculated by correlating the BOLD signal time series of the two participants within each dyad. This allowed for the creation of a dyad-level ISC matrix for each brain region. (B) The results of the first IS-RSA that linked categories of resilience and neural synchrony. Brain regions with significant differences for each contrast are outlined in black. (C) The calculation of resilience similarity in accordance with Anna Karenina model. (D) The results of the second IS-RSA that link resilience similarity and neural synchrony. Regions with significant associations between their neural synchrony and resilience similarity are outlined in black. False discovery rate (FDR) corrections were applied for all analyses. L, left hemisphere; R, right hemisphere.

Similar patterns were observed in the negative movie, but the group differences in ISC were found across a broader set of brain areas in all contrasts **(Fig.4B, Tab.S4)**. Specifically, higher ISCs of multiple brain regions that mainly anchored in the DAN, CN, and DMN were found in the {high, high} dyads than either {low, low} or {high, low} dyads. Moreover, ISCs of several brain regions including the visual cortex, inferior parietal lobe, lateral prefrontal cortex, and postcentral gyrus showed differences between {high, low} dyads and {low, low} dyads.

Collectively, our findings indicate greater neural synchrony in dyads of high-resilience pairs {high, high} compared to dyads of low-resilience pairs {low, low} or mixed pairs {high, low} during viewing both neutral and negative movies, suggesting that individuals with high resilience exhibit more consistent neural responses compared to their peers. In contrast, individuals with low resilience display more diverse neural responses among themselves.

To further investigate the relationship between brain synchrony and resilience, we performed a second IS-RSA utilizing resilience similarity to predict dyad-level ISC. The results revealed that resilience similarity significantly predicted neural synchrony in several brain regions, including the postcentral gyrus, frontal eye fields, and inferior parietal area, while participants were viewing the neutral movie (**Fig. 4D, Tab.S5**). Notably, many of these observed brain regions are core components of the DAN. For the negative movie, we found a positive association between resilience similarity and neural synchrony in areas of multiple brain networks including DMN (e.g., the inferior parietal lobe, posterior cingulate cortex, and precuneus), DAN (e.g., the frontal eye field, postcentral gyrus, superior parietal lobule), CN (e.g., the lateral prefrontal cortex and intraparietal sulcus), SMN, and VN (**Fig. 4D, Tab.S6**). Consistent with our primary results based on binarized resilience category, greater neural synchrony in individuals with higher resilience was found.

### Intolerance of Uncertainty modulates resilience-driven neural synchrony

Lastly, we tested the impact of IU on resilience-driven brain synchrony with the hypothesis that individuals with higher IU exhibit lower resilience-driven neural synchrony particularly in the DAN. As hypothesized, our findings indicated that increased IU inhibits the resilience-driven neural synchrony in areas associated with attention and perception (i.e., DAN subregions including the superior parietal lobe and postcentral gyrus), such that individuals with high IU and high resilience produce less neural synchrony in areas that are involved in attention engagement, visuospatial information processing, and motor-sensory function. the inhibition effect of IU on resilience-driven neural synchrony was also observed in the motor– and attention-related (e.g., primary motor sensory cortex, postcentral gyrus, and intraparietal sulcus) during the negative movie. We also found that the resilience-IU interaction significantly predicted neural synchrony during the neutral movie in the regions that functionally associated with motor-sensory (e.g., precentral gyrus), spatial attention (e.g., intraparietal sulcus and superior parietal lobule), face recognition (e.g., temporal areas) functions (**Fig. 5A, Tab.S7**). Our results indicated increased IU attenuated the positive association between neural synchrony and resilience similarity in these areas.

**Figure 5.**
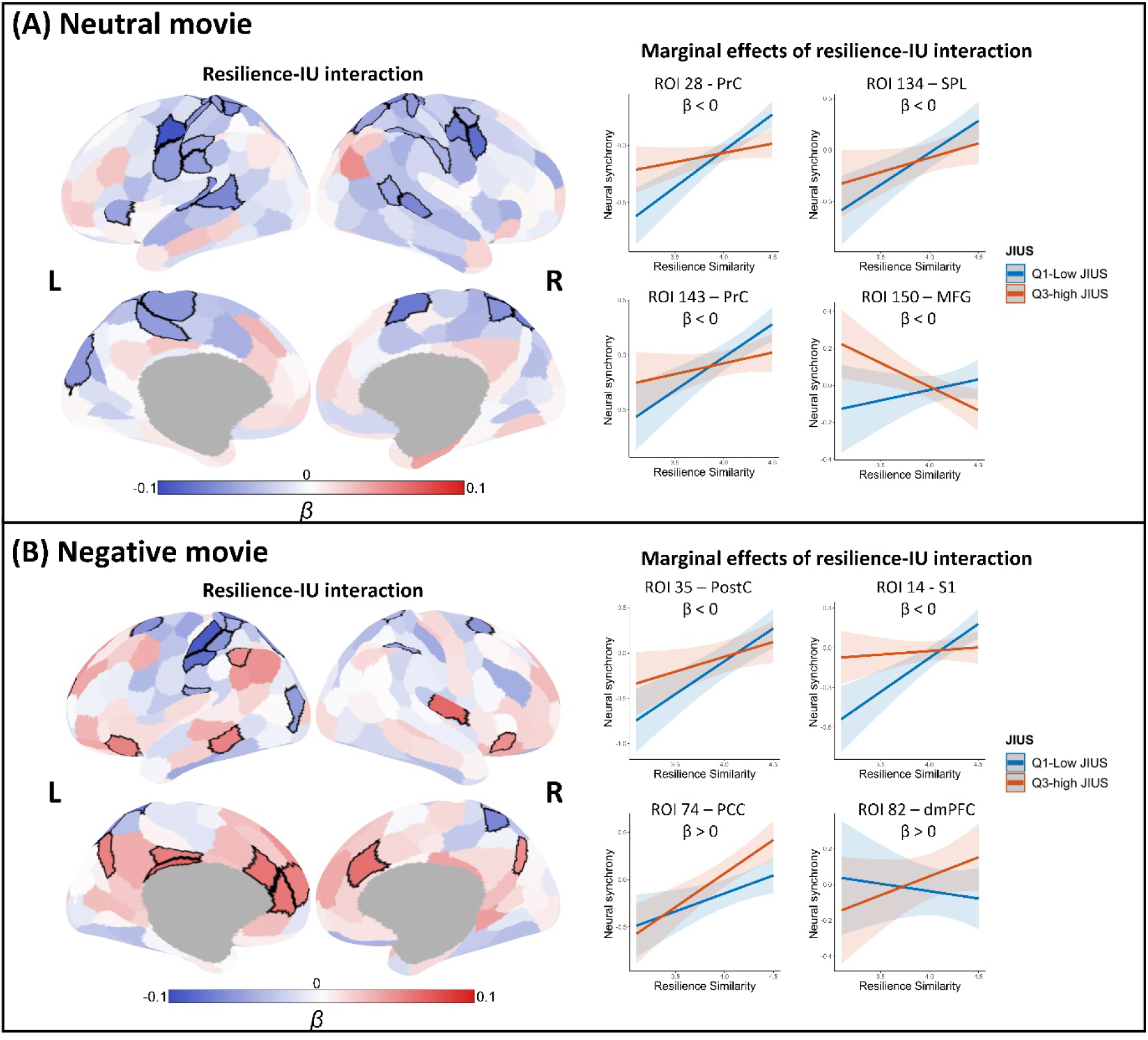
Modulation effects of intolerance of uncertainty (IU) on resilience-driven neural synchrony. (A) The results of the neutral movie. The brain maps illustrate that neural synchrony in brain areas, colored in blue and outlined in black, were dampened by the interaction between IU and resilience. Estimated marginal effects for neural synchrony by joint intolerance of uncertainty score (JIUS) indicated increased IU attenuated the resilience-driven neural synchrony. Four regions (bilateral precentral gyrus [ROI 28 and 143 – PrC], left superior parietal lobe [ROI 34 – SPL], and left middle frontal gyrus [ROI 150 – MFG]) that exhibited a large modulation effect of IU are presented as examples here. (B) The results of the negative movie. The brain maps illustrate that neural synchrony in brain areas, colored in blue and outlined in black, were dampened by the interaction between IU and resilience, and brain areas, colored in red and outlined in black, were enhanced by the IU-resilience interaction. Four regions (left postcentral gyrus [ROI 35 – PostC], left primary somatosensory cortex [ROI 14 – S1], left posterior cingulate cortex [ROI 74 – PCC], and left dorsomedial prefrontal cortex [ROI 82 – dmPFC]) that exhibited a large modulation effect of IU are presented as examples to demonstrate the estimated marginal effects. Q1 is the lower quartile, and Q3 is the upper quantile of JIUS. Shaded areas represent 95% CIs.

Intriguingly, we found that the resilience-IU interaction positively predicted ISC in the dorsomedial prefrontal cortex, ventral prefrontal cortex, posterior cingulate cortex, precuneus, inferior parietal lobe, and insula (**Fig. 5B, Tab.S8**) during the negative movie. This finding suggests that increased IU enhances the predictive ability of resilience similarity on neural synchrony of core regions of the DMN. These findings indicate that heightened IU intensified the resilience-driven neural synchrony in regions associated with affective functions when exposed to emotional stimuli. That is, resilient individuals with high IU shared more similar neural responses than their peers with low IU. Taken together, these findings indicate that increased IU weakens the resilience-driven neural synchrony of the DAN, but it enhances the resilience-driven neural synchrony of the DMN only when exposed to negative movie stimuli.

### Validation results

The validation analysis revealed consistent findings with our main results (**Fig.6)**. Therefore, the observed neural synchrony is driven by the similarities in participants’ resilience levels instead of the similarity in demographics including age, gender, and education.

**Figure 6.**
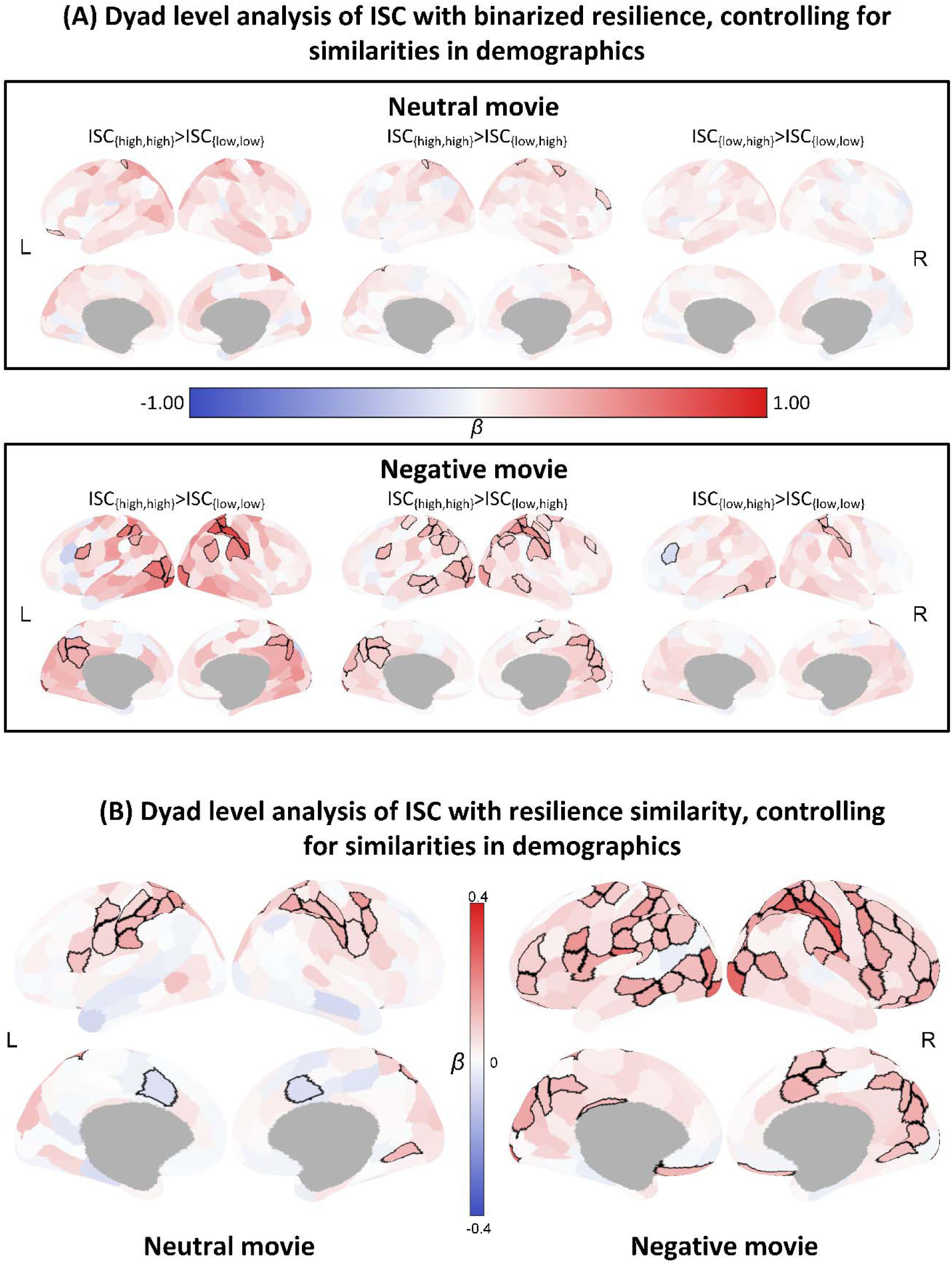
Results of IS-RSA validation. (A) Association of neural synchrony with binarized resilience while controlling for demographics including age, gender, and education. Similar patterns to our main findings were observed even after controlling for confounding variables. In the neutral condition, higher neural synchrony was observed in {high, high} dyads compared to {low, low} and {low, high} dyads. These patterns were even stronger in the negative condition. (B) Association of neural synchrony with resilience similarity while controlling for demographics including age, gender, and education. We identified similar brain regions that showed resilience-driven neural synchrony as our main findings. Regions with significant differences or associations are outlined in black. FDR corrections with a significance threshold of p < 0.05 were applied to address the issue of multiple comparisons.

## Discussion

Having a profound understanding of psychological resilience is vital to protect human beings from mental disorders. Nevertheless, to date, neuroimaging studies have not effectively investigated how resilience works as a complex system in an ecologically valid context, particularly considering the premise that resilience typically manifests after the encounter of negative events. Our findings revealed the convergent pattern of brain synchrony that is driven by resilience and uncovered the modulation effect of IU on resilience-driven neural synchrony. Our results establish an important contribution to understanding how individuals with varying levels of resilience perceive and process external stimuli, as reflected in neural synchrony, providing new neuropsychological evidence of why resilience should be considered as a dynamic and complex system rather than a static trait.

We discovered a convergent pattern of brain synchrony impacted by psychological resilience. Specifically, individuals with high resilience exhibit synchronized neural responses, while those with low resilience display divergent neural responses to external stimuli. This pattern concords with the AnnaK model, commonly employed to elucidate the vulnerability of multi-component systems(Finn et al., 2020; Zaneveld et al., 2017b). According to this model, system dysfunction occurs when deficits exist in any of the essential components. In the context of current study, the system refers to the resilience response which consists of multiple emotional stimulus processes. To maintain balanced mental states after exposure to negative stimuli, adaptive shaping of the entire processing chain, encompassing emotion perception, evaluation, integration, and regulation, is imperative (Foa et al., 2006; Gloria & Steinhardt, 2016). Resilient individuals sustain adaptability throughout each process, ultimately leading to successful responses to negative events. Conversely, dysfunction of any component may contribute to the risk of mental health disorders, and the risk may accumulate with an increasing number of failures within the process. Therefore, resilient individuals process the external stimuli in a highly similar manner, while low-resilience individuals show different brain responses as they may exhibit deficits in different processes. Our behavioral results, indicating a negative correlation between resilience and various emotion regulation difficulties, further support this assertion. Non-acceptance of emotional responses has been linked to hyperarousal symptoms and abnormal social-emotional evaluation(O’Bryan et al., 2015; Rudenstine et al., 2019), and lack of emotional clarity has been associated with impairment in emotion recognition and self-referential processing(Faulkner et al., 2020; Gratz & Roemer, 2004). Our findings thus elucidate that resilient individuals can effectively maintain their mental health(Ungar & Theron, 2020), whereas individuals with low resilience exhibit more emotion regulation difficulties(Kim et al., 2021; Poole et al., 2017) and are more susceptible to various mental disorders(Bhatnagar, 2021; Herrman et al., 2011).

Resilience-driven brain synchrony was observed in regions anchored in the DAN. During naturalistic movie-watching, top-down attention enhances focus from a specific perspective, aids in gathering relevant information, and enables the spread of information across brain networks(Regev et al., 2019). Consequently, the brain synchrony in the DAN may reflect the aligned neural encoding of external stimuli(Nummenmaa et al., 2012), implying a shared perspective among participants during movie watching. As the emotional intensity of stimuli increased, during the negative movie clip, the resilience-driven brain synchrony extended to other brain networks including the DMN and CN. Empirical evidence suggested that the high neural similarity of DMN and CN signifies shared narrative interpretation among audiences(Nguyen et al., 2019). The DMN is reported as a key network in social cognition, with one of its core functions being the respond to the actions of other social agents and characterization of interpersonal interactions(Yeshurun et al., 2021). Previous studies have found that the increased DMN functional coupling between speaker and listener predicts a more similar understanding of the story between them(Silbert et al., 2014). The CN is wildly implicated in cognitive control and emotion regulation, fundamental for facilitating the understanding of events(Morawetz et al., 2020; Niendam et al., 2012). High brain synchrony within these two networks suggests a similarity in their interpretation of the movie among resilient individuals, possibly stemming from their similarity in emotional regulation and evaluation when processing emotional stimuli. Based on these findings, we speculate that resilience reflects the overall adaptability of emotional functions, requiring coordination amongst multiple networks, primarily including the DAN, DMN, and CN.

The current findings align with previous research indicating that resilience involves multiple brain networks and is closely intertwined with emotional processing(Li et al., 2024). Building upon these neuroimaging discoveries and recognizing the intricate nature of emotional processes to negative events, it is important to consider resilience as an ongoing and dynamic neural process that encompasses dimensions of emotion, cognition, and attention(Scherer & Moors, 2019). This dynamicity characteristic is well manifested while participants engage in processing naturalistic stimuli that unfold over time, thus providing a methodological advantage of uncovering neural correlates associated with resilience that are dismissed using traditional task-based design. While our study did not specifically focus on emotional recognition, it extends previous hypotheses about emotions, such as the constructionist approach(Lindquist et al., 2012; Xu et al., 2021), to naturalistic movies, which posits that multiple brain networks critical for social evaluation, emotion regulation, and attention orientation when encoding naturalistic movies with emotional valence(Kringelbach et al., 2023; Nummenmaa et al., 2023). Therefore, future studies should incorporate more naturalistic movies to test hypotheses and models of emotions, aiming to examine the generalizability of existing models to more dynamic and naturalistic stimuli.

Additionally, we made a noteworthy discovery that IU plays a role in modulating resilience-driven brain synchrony. Consistent evidence from empirical studies across diverse samples demonstrated a negative association between IU and resilience(Di Trani et al., 2021; Panzeri et al., 2021; T. Wang et al., 2023). Here, we found increased IU led to a reduction in resilience-driven brain synchrony, primarily in brain regions belong to the DAN. Prior research has indicated a negative association between IU and attention inhibition, which is primarily governed by the DAN(Morriss & McSorley, 2019). This suggests that weak goal-oriented attention processing, caused by high IU, may be linked with a lack of focused attention to external stimuli and highly variable activity in the DAN, leading to decreased resilience-driven brain synchrony. Alternatively, IU, as a risk factor for mental disorders, is associated with attention dysfunction(Gramszlo et al., 2018). Higher IU is linked to abnormal attentional functions, thus contributing to divergent neural activity within the DAN. Consequently, the inhibitory modulation of IU aligns with the AnnaK model, which emphasizes the importance of intact components in a system to maintain proper functioning. In other words, IU dampens resilience by impairing the attention functions critical for gating external information and gathering emotional cues. Moreover, increased IU leads to a biased prediction error, which is critical in updating anticipation and reorienting attention(Nelson et al., 2016; Sinclair et al., 2021). Meanwhile, the prediction error largely relies on the dopamine signaling in the nervous system(Papalini et al., 2020). Previous research also demonstrated the role of dopamine in supporting coupling of DAN with other large scale networks such as the DMN(Dang et al., 2012). Thus, the pharmacological intervention that targets dopamine might be a promising avenue to modulate individuals’ IU levels, thereby enhancing resilience and fostering more adaptive behaviors (e.g., attention engagement and emotion regulation), reducing the risk of psychiatric disorders.

We also found that increased IU enhanced the resilience-driven brain synchrony within the DMN areas only during watching negative movie. The DMN exhibited greater brain synchrony in individuals who were both highly resilient and uncertainty-intolerant. Considering the role of DMN in generating shared insights into events(Yeshurun et al., 2021), these findings may suggest that increased IU promotes the formation of a coherent, clear, and certain interpretation and understanding of emotional narratives in resilient individuals. This could allow them to avoid negative feelings stemming from the uncertainty associated with novel emotional stimuli.

Several limitations should be noted when interpreting our findings. Firstly, we relied on the self-report questionnaire to measure individuals’ psychological resilience, which may not fully represent its dynamic nature over an extended period, despite it somehow reflecting the general level of resilience. Therefore, conducting future prospective longitudinal studies would be beneficial in addressing this limitation and gaining a more comprehensive understanding of how resilience evolves over time. Secondly, the focus on young adults in our study may restrict the generalizability of the findings to broader populations. To overcome this limitation, future research should consider including more diverse samples, encompassing individuals from different age groups, such as the clinical sample and older adult samples. Thirdly, neutral and negative stimuli were exclusively employed in the current study. Although resilience typically functions when exposed to negative stimuli, previous works have shown that resilience could modulate responses to positive stimuli processing(Ohad & Yeshurun, 2023; Rademacher et al., 2023; Thoern et al., 2016). Thus, incorporating positive stimuli in future studies would provide a more balanced perspective and enable us to examine how resilience influences neural responses to the variety of emotional valences and arousal levels.

In conclusion, the current study revealed that resilience plays a key role in eliciting neural synchrony among individuals during movie viewing. This resilience-driven brain synchrony aligns with the AnnaK model, wherein resilient individuals share similar neural responses, in a synchronized manner with a coherency among multiple brain networks involving attention, cognitive control, and social cognition, while each low-resilience individual exhibits divergent neural activity and lower synchrony among themselves. Moreover, we propose the modulation effect of IU on resilience-driven synchrony, which highlights the role of IU in the resilience system and its associated brain responses. This knowledge deepens our understanding of resilience as a complex system and may potentially inform clinical and therapeutic interventions aimed at individualized approaches in fostering resilience, moving towards the precision psychiatry framework to improve mental health outcomes among the clinical population.

## Acknowledgments

We would like to acknowledge the help we received from radiographers and MR physicists at the 7T MR centre at NTNU. We thank our participants for their time and effort during the experiment, Jørgen Østmo-Sæter Olsnes for his help during experimental setup, and Stian Framvik for his help with the data acquisition.

## Data, Materials, and Software Availability

Analysis codes are available at OSF | Video watching and resilience (https://osf.io/cgjkf/). The raw data will be available from the first author upon request.

