## Supplemental Information for "Resilience-driven neural synchrony during naturalistic movie watching"

**This PDF file includes:**

Supporting information

Figures S1

Tables S1 to S8

SI References

**Supporting Information**

**Pilot experiment for selecting movie clips**

Four movie clips, each lasting about 8 minutes, were sourced from the internet and previous studies (Taylor et al., 2017). The first clip was a documentary showcasing various animals accompanied by soft background music entitled “*Animals*”. The second clip, titled “*Pottery”*, documents the processing of two women making pottery. The third clip is a story of a woman desperately attempting to maintain her balance on a slope of a dam to prevent herself from plummeting into the abyss below entitled “*Curve*”. The fourth is a clip from the famous black-white movie “*Bang! You’re Dead”* from 1961*(directed by Alfred Hitchcock)*, tells a thrilling story that A 5-year-old boy finds his uncle's revolver, partially loads it with bullets, and plays with it in public, unaware of its deadly power. Based on the content and previous studies, the movie *Animals* and *Pottery* are considered as neutral clips, while *Curve* and *Bang! You’re* dead are considered as negative ones.

To examine the emotional property, including the emotional valence and arousal of the movie clips, a pilot experiment was conducted. We recruited 24 participants (12 males, mean age =25.5 ±3.96 yrs) from the local community to rate these four movies. A MATLAB-based software for Continuous Affect Rating and Media Annotation (CARMA, v14.10) was used to play the movies and to acquire continuous ratings on emotional intensity ranging from 0 to 100 (Girard, 2014). During the movie watching, participants were asked to indicate their emotional intensity they experienced by adjusting a cursor on a scroll in the bottom of the screen. The rating feedback was sampled at the rate of 1 Hz (i.e., sample the position of the cursor one time per second). Following watching each movie, they provided ratings on the intensity of various emotions (e.g., happiness, anger, stress, satisfaction, etc.) as well as their perception of emotional valence and arousal induced by the movies.

To avoid the spillover effect, the neutral movie clips were presented first, followed by the negative movies. The sequence of movie clips within the same condition was counterbalanced across participants (see **Fig. S1a**). None of the participants had watched any of these movie clips before the experiment.

The results of continuous emotion-intensity feedback are presented in **Fig. S1.** For the neutral movie clips, *Pottery* triggered lower emotional intensity than *Animals* (*t* = -16.304, *p* < 0.001, *d* = 3.033). For negative movies, participants experienced higher emotion intensity in *Curve* rather than *Bang! You’re dead* (*t =* 45.377*, p <* 0.001, *d* = 2.911). In the current sample, the average-rating consistency intraclass correlation coefficient is 0.823 (*Animals*), 0.818 (*Pottery*),0.976 (*Curve*), and 0.969 (*Bang! You’re Dead*), indicating good reliability of the continuous rating for all movie clips.

The results of the emotion rating following movie watching are presented in **Fig. S5**. Participants predominantly reported feeling happy, interested, relaxed, and satisfied while watching both neutral movies. The overall intensity of these major emotions in *Pottery* is lower than in *Animals* indicating *Animals* induced stronger emotion experience. In the comparison between two neutral movies, *Animals* and *Pottery*, significantly higher emotional valence (mean*_Animals_ =* 6.917*,* mean*_Pottery_ =* 5.875*, t =* 3.824*, p <* 0.001, *d* =0.781) and arousal (mean*_Animals_ =* 4.083*,* mean*_Pottery_ =* 2.958*, t =* 2.944*, p =* 0.007, *d* = 0.601 ) were observed in *Animals* compared to *Pottery.* In the negative movie clips, participants reported high level of stress, fear, and pain during the *Curve*, while *Bang! You’re Dead* elicited feelings of stress, fear, and anger in participants. Moreover, participants reported overall higher valence and lower arousal for the negative movies compared to neutral ones. In the comparison between two negative movies, *Curve* and *Bang! You’re Dead,* a marginally significant difference was observed in emotional valence (mean*_Curve_ =* 3.125*,* mean*_Bang! You’re Dead_ =* 3.750*, t =* -2.044*, p =* 0.051, *d* = 0.417) while no differences in arousal (mean*_Curve_ =* 6.542*,* mean*_Bang! You’re Dead_ =* 6.125*, t =* 1.551*, p =* 0.134, *d* =0.317) were obtained. Taken together, the movies we considered as neutral (i.e., *Animals* and *Pottery*) were distinctively different from the negative movie clips (i.e., *Curve* and *Bang!You’re Dead*). Compared with negative movies, neutral movies induced fewer negative emotions, higher emotional valence, lower emotional arousal, and lower continuous experienced emotional intensity during the viewing.

Most previous fMRI experiments used nature documentaries similar to *Animals*, as a neutral condition(Camacho et al., 2019; Chen et al., 2020; van Baar et al., 2021). In our experiment, however, we opted to use *Pottery* as the neutral stimulus for the following reasons: 1) Based on the results of the emotion rating, *Animals* received more positive ratings on emotional valence and evoked greater emotional arousal and intensity in the audience compared with *Pottery*, suggesting that it leans more towards being a positive stimulus instead of a neutral one.2) In *Animals*, soft background music accompanies the scenes, whereas in *Pottery*, only the real sounds from the scenes, such as the characters' actions, are present without any additional music making this clip similar to negative stimuli and closer to real life. 3) Finally, to have more comparison with the negative movie clips which included human faces, the Animals did not include any human faces and thus the pottery was chosen instead.

For negative movies, although *Bang! You're Dead* has been applied as the negative movie to induce tension and stress in previous studies (Taylor et al., 2017), we used *Curve* rather than *Bang! You’re Dead* with the following reasons: 1). Based on the results, *Curve* triggered stronger and more types of negative emotions than *Bang! You’re Dead*, 2) *Bang! You’re Dead is* predominantly spoken in English, and this language preference may impact the information processing of non-native English speakers. On the other hand, *Curve* is a movie without dialogue, making it more accessible and compatible to speakers of other languages.

Moreover, the selected movies (i.e., *Pottery* and *Curve*) shared the following characteristics to control for potential confounding effects: 1) No more than one human character is present in each clip at any given time; 2) all the depicted characters are females; 3) Absence of dialogue, word descriptions, and subtitles. 4) Both movies are presented in color.

**Table S1. The website for each movie clips.**

| **Movie clips** | **Website** |
| --- | --- |
| *Animals* | <https://www.youtube.com/watch?v=7IBzc3pajzY&list=PLJtPH8t_isfXtrfesaF0n5CnneIqPfSTt&index=15> |
| *Pottery* | <https://www.youtube.com/watch?v=Qi4JQEBW6Kw> |
| *Curve* | <https://www.youtube.com/watch?v=2dD3Fawk4y0> |
| *Bang!You’re Dead* | - |

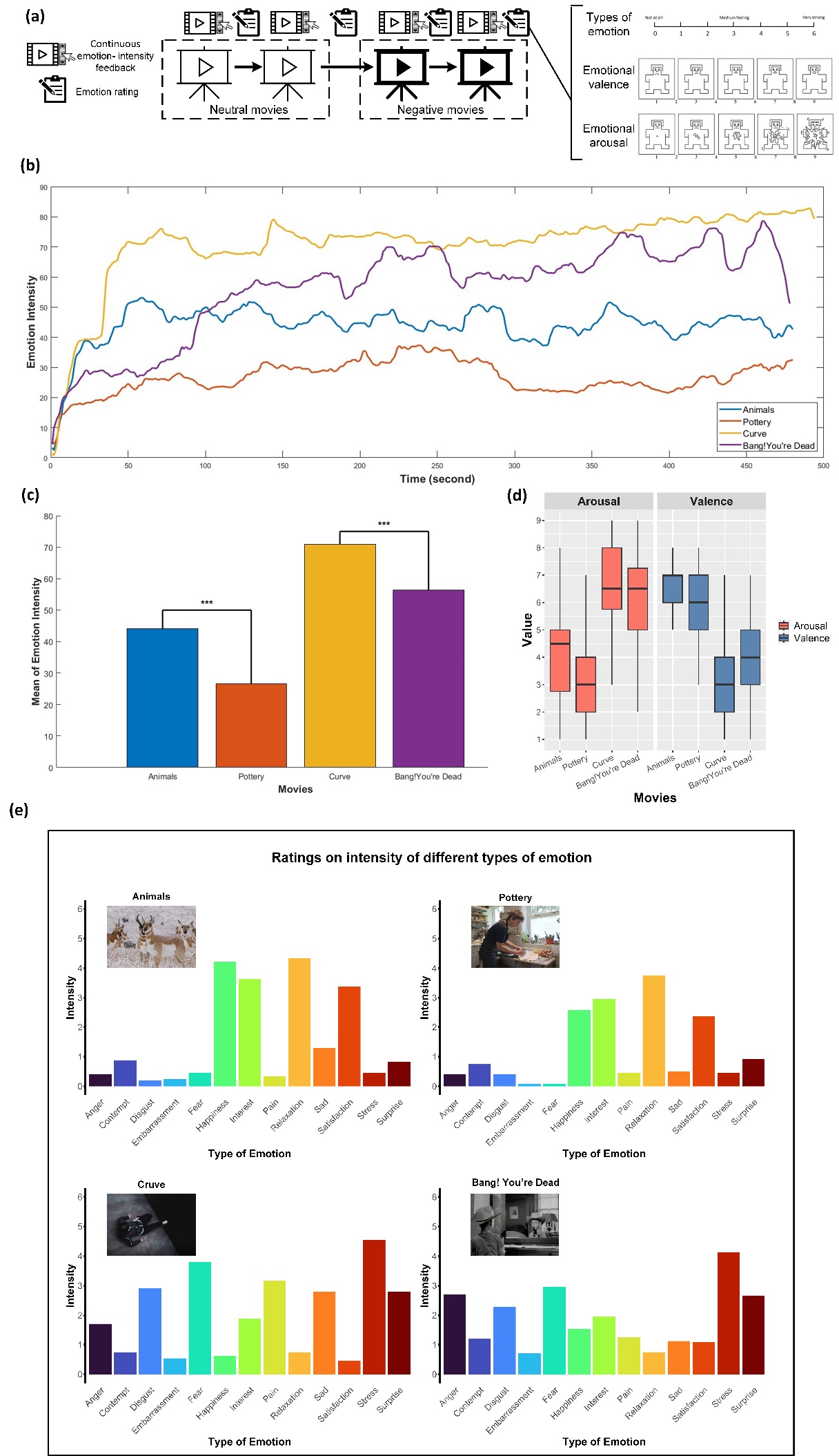

**Fig. S1. Workflow and results of the movie-rating. (a)** Participants were instructed to watch 4 movie clips (2 neutral and 2 negative movies). During the viewing process, participants provided continuous feedback regarding the emotional intensity they experienced. After watching each movie, they rated the intensity of various emotions, as well as emotional valence and arousal. **(b)** Changes in emotion intensity induced by the movies over the course of movie clips indicating lower intensity overall for the *Pottery* and highest for the *Curve*. Panel **(c)** represents the bar plot of the mean of the emotion intensity for each movie, indicating higher emotional intensity for the *Curve* and *Animals* compared to the *Bang! You’re dead* and *Pottery*, respectively. Panel **(d)** represents the results of ratings on emotional valence and arousal. The lower and upper edges of the box plot represent the first quartile and third quartile, respectively. The line inside the box represents the median. Panel **(e)** indicates the results of ratings on various emotions experienced during movie-watching. ^***^: *p* < 0.001

**Table S2. Participant-level ISC of subcortical areas.**

| Subcortical region | ISC (Mean ± SD) | | *t*-value |
| --- | --- | --- | --- |
|  | Neutral movie | Negative movie |  |
| Hippocampus (R) | 0.126 ± 0.159 | 0.257 ± 0.192 | -4.826^*^ |
| Amygdala (R) | 0.155 ± 0.138 | 0.283 ± 0.151 | -6.248^*^ |
| Posterior Thalamus (R) | 0.179 ± 0.162 | 0.269 ± 0.186 | -3.894 |
| Anterior Thalamus (R) | 0.208 ± 0.141 | 0.274 ± 0.152 | -3.086 |
| Nucleus accumbens (R) | 0.151 ± 0.139 | 0.173 ± 0.160 | -0.915 |
| Globus pallidus (R) | 0.071 ± 0.113 | 0.213 ± 0.218 | -4.848^*^ |
| Putamen (R) | 0.036 ± 0.151 | 0.211 ± 0.186 | -5.626^*^ |
| Caudate nucleus (R) | 0.058 ± 0.127 | 0.218 ± 0.202 | -5.752^*^ |
| Hippocampus (L) | 0.148 ± 0.186 | 0.239 ± 0.197 | -3.402 |
| Amygdala (L) | 0.141 ± 0.127 | 0.255 ± 0.154 | -5.295^*^ |
| Posterior Thalamus (L) | 0.150 ± 0.178 | 0.264 ± 0.191 | -4.167 |
| Anterior Thalamus (L) | 0.179 ± 0.137 | 0.251 ± 0.168 | -3.402 |
| Nucleus accumbens (L) | 0.167 ± 0.142 | 0.179 ± 0.156 | -0.619 |
| Globus pallidus (L) | 0.059 ± 0.116 | 0.208 ± 0.203 | -5.265^*^ |
| Putamen (L) | 0.032 ± 0.135 | 0.200 ± 0.178 | -5.815^*^ |
| Caudate nucleus (L) | 0.082 ± 0.129 | 0.230 ± 0.183 | -5.757^*^ |

FDR correction was applied to correct for multiple comparisons. All the reported *p*-values are two-tailed. SD: standard deviation; R: right; L: left; ^*^: corrected *p* < 0.05.

**Table S3. Dyad-level results that relate binarized resilience with ISC during neutral movie.**

|  | Region | β | | Standard Error | | Corrected  *p*-value | Network |
| --- | --- | --- | --- | --- | --- | --- | --- |
|  | **Contrast: ISC_{high, high}_ > ISC_{low, low}_** | | | | | | |
|  | ***Significant cortical regions*** | | | | | | |
|  | No. 93 PFCv_2 | 0.231 | | | 0.085 | 0.025 | DMN |
|  | No. 17 SomMotA_5 | 0.319 | | | 0.124 | 0.028 | SMN |
|  | No. 139 PostC_3 | 0.455 | | | 0.180 | 0.028 | DAN |
|  | No. 39 FEF_1 | 0.439 | | | 0.184 | 0.044 | DAN |
|  | ***Subcortical regions*** |  | | |  |  |  |
|  | Hippocampus (R) | 0.071 | | | 0.111 | 0.609 | SUB |
|  | Amygdala (R) | -0.035 | | | 0.106 | 0.823 | SUB |
|  | Posterior Thalamus (R) | 0.000 | | | 0.128 | 0.997 | SUB |
|  | Anterior Thalamus (R) | 0.146 | | | 0.124 | 0.322 | SUB |
|  | Nucleus accumbens (R) | -0.106 | | | 0.105 | 0.387 | SUB |
|  | Globus pallidus (R) | 0.032 | | | 0.066 | 0.706 | SUB |
|  | Putamen (R) | 0.010 | | | 0.079 | 0.938 | SUB |
|  | Caudate nucleus (R) | -0.055 | | | 0.073 | 0.545 | SUB |
|  | Hippocampus (L) | 0.024 | | | 0.135 | 0.906 | SUB |
|  | Amygdala (L) | 0.002 | | | 0.098 | 0.992 | SUB |
|  | Posterior Thalamus (L) | 0.039 | | | 0.125 | 0.832 | SUB |
|  | Anterior Thalamus (L) | 0.121 | | | 0.109 | 0.342 | SUB |
|  | Nucleus accumbens (L) | -0.078 | | | 0.116 | 0.602 | SUB |
|  | Globus pallidus (L) | -0.008 | | | 0.066 | 0.939 | SUB |
|  | Putamen (L) | 0.032 | | | 0.069 | 0.720 | SUB |
|  | Caudate nucleus (L) | -0.057 | | | 0.079 | 0.567 | SUB |
|  | **Contrast: ISC_{high, high}_ > ISC_{low, high}_** | | | | | | |
| ***Significant cortical regions*** | | | | | | | |
| No. 140 PostC_4 | | | 0.362 | | 0.118 | 0.010 | DAN |
| No. 153 PFCl_1 | | | 0.190 | | 0.067 | 0.019 | CN |
| No. 17 SomMotA_5 | | | 0.206 | | 0.077 | 0.025 | SMN |
| No. 38 PostC_4 | | | 0.290 | | 0.114 | 0.028 | DAN |
| No. 114 SomMotA_2 | | | 0.178 | | 0.070 | 0.028 | SMN |
| No. 139 PostC_3 | | | 0.249 | | 0.099 | 0.028 | DAN |
| No. 141 FEF_1 | | | 0.277 | | 0.111 | 0.028 | DAN |
| ***Subcortical regions*** | | |  | |  |  |  |
| Hippocampus (R) | | | -0.030 | | 0.072 | 0.759 | SUB |
| Amygdala (R) | | | -0.042 | | 0.071 | 0.627 | SUB |
| Posterior Thalamus (R) | | | -0.013 | | 0.078 | 0.913 | SUB |
| Anterior Thalamus (R) | | | 0.115 | | 0.077 | 0.206 | SUB |
| Nucleus accumbens (R) | | | -0.027 | | 0.070 | 0.777 | SUB |
| Globus pallidus (R) | | | 0.067 | | 0.058 | 0.326 | SUB |
| Putamen (R) | | | 0.036 | | 0.061 | 0.645 | SUB |
| Caudate nucleus (R) | | | -0.006 | | 0.060 | 0.948 | SUB |
| Hippocampus (L) | | | -0.053 | | 0.081 | 0.607 | SUB |
| Amygdala (L) | | | -0.042 | | 0.068 | 0.613 | SUB |
| Posterior Thalamus (L) | | | -0.008 | | 0.077 | 0.941 | SUB |
| Anterior Thalamus (L) | | | 0.078 | | 0.071 | 0.349 | SUB |
| Nucleus accumbens (L) | | | -0.060 | | 0.074 | 0.501 | SUB |
| Globus pallidus (L) | | | -0.007 | | 0.058 | 0.939 | SUB |
| Putamen (L) | | | 0.038 | | 0.059 | 0.607 | SUB |
| Caudate nucleus (L) | | | -0.039 | | 0.062 | 0.609 | SUB |
| **Contrast: ISC_{low, high}_ > ISC_{low, low}_** | | | | | | | |
| ***Subcortical regions*** | |  | | |  |  |  |
| Hippocampus (R) | | 0.101 | | | 0.072 | 0.214 | SUB |
| Amygdala (R) | | 0.008 | | | 0.071 | 0.939 | SUB |
| Posterior Thalamus (R) | | 0.014 | | | 0.078 | 0.906 | SUB |
| Anterior Thalamus (R) | | 0.031 | | | 0.077 | 0.763 | SUB |
| Nucleus accumbens (R) | | -0.079 | | | 0.070 | 0.326 | SUB |
| Globus pallidus (R) | | -0.035 | | | 0.058 | 0.609 | SUB |
| Putamen (R) | | -0.025 | | | 0.061 | 0.752 | SUB |
| Caudate nucleus (R) | | -0.049 | | | 0.060 | 0.480 | SUB |
| Hippocampus (L) | | 0.077 | | | 0.081 | 0.411 | SUB |
| Amygdala (L) | | 0.045 | | | 0.068 | 0.601 | SUB |
| Posterior Thalamus (L) | | 0.047 | | | 0.077 | 0.613 | SUB |
| Anterior Thalamus (L) | | 0.043 | | | 0.071 | 0.617 | SUB |
| Nucleus accumbens (L) | | -0.017 | | | 0.074 | 0.872 | SUB |
| Globus pallidus (L) | | -0.001 | | | 0.058 | 0.993 | SUB |
| Putamen (L) | | -0.006 | | | 0.059 | 0.939 | SUB |
| Caudate nucleus (L) | | -0.018 | | | 0.062 | 0.836 | SUB |

The index of cortical regions is taken from Schafer 200 atlas. FDR correction was applied to correct for multiple comparisons. All the reported *p*-values are two-tailed. R: right; L: left; DAN: dorsal attention network; CN: control network; DMN: default mode network; SMN: somatomotor network; SUB: subcortical network.

**Table S4. Dyad-level results that relate binarized resilience with ISC during negative movie.**

|  | Region | β | SE | | Corrected  *p*-value | Network |
| --- | --- | --- | --- | --- | --- | --- |
|  | **Contrast: ISC_{high, high}_ > ISC_{low, low}_** | | | | | |
|  | ***Significant cortical region*** | | | | | |
|  | No. 135 SPL_3 | 0.708 | | 0.235 | 0.001 | DAN |
|  | No. 138 PostC_2 | 0.848 | | 0.290 | 0.002 | DAN |
|  | No. 2 ExStr_2 | 0.728 | | 0.254 | 0.002 | VN |
|  | No. 137 Post_C1 | 0.771 | | 0.275 | 0.003 | DAN |
|  | No. 140 Post_C4 | 0.709 | | 0.263 | 0.006 | DAN |
|  | No. 117 SomMotA_5 | 0.725 | | 0.272 | 0.006 | SMN |
|  | No. 104 ExStr_2 | 0.618 | | 0.245 | 0.011 | VN |
|  | No. 79 pCunPCC_3 | 0.434 | | 0.184 | 0.019 | DMN |
|  | No. 64 PFCl_2 | 0.498 | | 0.215 | 0.020 | CN |
|  | No. 68 IPL_1 | 0.408 | | 0.177 | 0.021 | CN |
|  | No. 182 pCun_2 | 0.448 | | 0.196 | 0.021 | CN |
|  | No. 181 pCun_1 | 0.509 | | 0.226 | 0.024 | CN |
|  | No. 36 PostC_2 | 0.588 | | 0.263 | 0.024 | DAN |
|  | No. 118 SomMotA_6 | 0.252 | | 0.113 | 0.024 | SMN |
|  | No. 184 IPL_1 | 0.448 | | 0.202 | 0.025 | DMN |
|  | No. 60 IPS_3 | 0.487 | | 0.224 | 0.027 | CN |
|  | No. 166 IPS_2 | 0.471 | | 0.217 | 0.028 | CN |
|  | No. 172 Temp_2 | 0.399 | | 0.187 | 0.029 | CN |
|  | No. 31 ParOcc_1 | 0.609 | | 0.289 | 0.033 | DAN |
|  | No. 75 IPL_1 | 0.386 | | 0.186 | 0.035 | DMN |
|  | No. 77 pCunPCC_1 | 0.349 | | 0.168 | 0.035 | DMN |
|  | No. 142 ParOper_1 | 0.550 | | 0.268 | 0.035 | VAN |
|  | No. 72 pCun_1 | 0.430 | | 0.211 | 0.035 | CN |
|  | No. 115 SomMotA_3 | 0.320 | | 0.157 | 0.035 | SMN |
|  | No. 5 ExStr_4 | 0.532 | | 0.263 | 0.036 | VN |
|  | No. 16 SomMotA_4 | 0.279 | | 0.139 | 0.037 | SMN |
|  | No. 186 pCunPCC_1 | 0.384 | | 0.192 | 0.038 | DMN |
|  | No. 39 EFE_1 | 0.336 | | 0.170 | 0.041 | DAN |
|  | No. 37 PostC_3 | 0.565 | | 0.289 | 0.043 | DAN |
|  | No. 67 Temp_1 | 0.334 | | 0.171 | 0.043 | CN |
|  | No. 120 SomMotA_8 | 0.491 | | 0.254 | 0.046 | SMN |
|  | No. 113 SomMotA_1 | 0.332 | | 0.173 | 0.047 | SMN |
|  | No. 86 Temp_4 | 0.402 | | 0.201 | 0.047 | DMN |
|  | No. 173 IPL_1 | 0.400 | | 0.210 | 0.048 | CN |
|  | ***Subcortical region*** | | | | | |
|  | Hippocampus (R) | 0.236 | | 0.192 | 0.228 | SUB |
|  | Amygdala (R) | 0.060 | | 0.164 | 0.753 | SUB |
|  | Posterior Thalamus (R) | -0.128 | | 0.189 | 0.520 | SUB |
|  | Anterior Thalamus (R) | -0.066 | | 0.163 | 0.722 | SUB |
|  | Nucleus accumbens (R) | -0.021 | | 0.137 | 0.902 | SUB |
|  | Globus pallidus (R) | 0.103 | | 0.189 | 0.621 | SUB |
|  | Putamen (R) | -0.199 | | 0.170 | 0.252 | SUB |
|  | Caudate nucleus (R) | -0.087 | | 0.182 | 0.671 | SUB |
|  | Hippocampus (L) | 0.154 | | 0.188 | 0.421 | SUB |
|  | Amygdala (L) | 0.201 | | 0.150 | 0.189 | SUB |
|  | Posterior Thalamus (L) | -0.076 | | 0.190 | 0.726 | SUB |
|  | Anterior Thalamus (L) | -0.030 | | 0.165 | 0.889 | SUB |
|  | Nucleus accumbens (L) | -0.194 | | 0.125 | 0.119 | SUB |
|  | Globus pallidus (L) | -0.050 | | 0.184 | 0.823 | SUB |
|  | Putamen (L) | -0.161 | | 0.157 | 0.320 | SUB |
|  | Caudate nucleus (L) | -0.163 | | 0.172 | 0.344 | SUB |
|  | **Contrast: ISC_{high, high}_ > ISC_{low, high}_** | | | | | |
| ***Significant cortical region*** | | | | | | |
| No. 135 SPL_3 | | 0.476 | | 0.123 | <0.001 | DAN |
| No. 2 ExStr_2 | | 0.448 | | 0.131 | <0.001 | VN |
| No. 138 PostC_2 | | 0.506 | | 0.147 | <0.001 | DAN |
| No. 182 pCun_2 | | 0.371 | | 0.106 | <0.001 | CN |
| No. 104 ExStr_3 | | 0.422 | | 0.127 | <0.001 | VN |
| No. 137 PostC_1 | | 0.459 | | 0.140 | <0.001 | DAN |
| No. 140 PostC_4 | | 0.432 | | 0.135 | 0.001 | DAN |
| No. 117 SomMotA_5 | | 0.441 | | 0.139 | 0.001 | SMN |
| No. 79 pCunPCC_3 | | 0.315 | | 0.101 | 0.001 | DMN |
| No. 64 PFCl_2 | | 0.335 | | 0.114 | 0.002 | CN |
| No. 165 IPS_1 | | 0.272 | | 0.094 | 0.002 | CN |
| No. 166 IPS_2 | | 0.314 | | 0.115 | 0.005 | CN |
| No. 86 Temp_4 | | 0.292 | | 0.112 | 0.008 | DMN |
| No. 115 SomMotA_3 | | 0.233 | | 0.090 | 0.008 | SMN |
| No. 36 PostC_2 | | 0.349 | | 0.135 | 0.009 | DAN |
| No. 75 IPL_1 | | 0.256 | | 0.102 | 0.011 | CN |
| No. 181 pCun_1 | | 0.298 | | 0.119 | 0.011 | CN |
| No. 12 ExStrSup_2 | | 0.301 | | 0.124 | 0.016 | CN |
| No. 68 IPL_1 | | 0.237 | | 0.098 | 0.016 | CN |
| No. 118 SomMotA_6 | | 0.175 | | 0.073 | 0.016 | SMN |
| No. 60 IPS_3 | | 0.280 | | 0.118 | 0.019 | CN |
| No. 32 SPL_1 | | 0.276 | | 0.117 | 0.019 | DAN |
| No. 58 IPS_1 | | 0.180 | | 0.077 | 0.020 | CN |
| No. 173 IPL_1 | | 0.261 | | 0.112 | 0.020 | CN |
| No. 5 ExStr_4 | | 0.309 | | 0.135 | 0.021 | VN |
| No. 31 ParOcc_1 | | 0.335 | | 0.147 | 0.021 | DAN |
| No. 67 Temp_1 | | 0.217 | | 0.096 | 0.021 | CN |
| No. 172 Temp_2 | | 0.234 | | 0.102 | 0.021 | CN |
| No. 175 PFCld_1 | | 0.186 | | 0.082 | 0.021 | CN |
| No. 16 SomMotA_4 | | 0.183 | | 0.082 | 0.024 | SMN |
| No. 133 SPL_1 | | 0.313 | | 0.142 | 0.025 | DAN |
| No. 109 StriCal_1 | | 0.289 | | 0.132 | 0.026 | VN |
| No. 120 SomMotA_8 | | 0.287 | | 0.131 | 0.027 | SMN |
| No. 113 SomMotA_1 | | 0.208 | | 0.096 | 0.027 | SMN |
| No. 111 ExStrSup_2 | | 0.272 | | 0.126 | 0.028 | VN |
| No. 37 PostC_3 | | 0.313 | | 0.147 | 0.031 | DAN |
| No. 39 FEF_1 | | 0.195 | | 0.095 | 0.035 | DAN |
| No. 42 FrOper_1 | | 0.204 | | 0.099 | 0.035 | VAN |
| No. 59 IPS_2 | | 0.184 | | 0.090 | 0.035 | CN |
| No. 72 pCun_1 | | 0.230 | | 0.112 | 0.035 | CN |
| No. 77 pCunPCC_1 | | 0.194 | | 0.094 | 0.035 | DMN |
| No. 136 SPL_4 | | 0.273 | | 0.133 | 0.035 | DAN |
| No. 142 ParOper_1 | | 0.283 | | 0.138 | 0.035 | VAN |
| No. 184 IPL_1 | | 0.224 | | 0.109 | 0.035 | DMN |
| No. 176 PFCld_2 | | 0.189 | | 0.093 | 0.037 | CN |
| No. 143 PrC_1 | | 0.211 | | 0.105 | 0.038 | VAN |
| No. 26 S2_3 | | 0.237 | | 0.121 | 0.043 | SMN |
| No. 195 Rsp_1 | | 0.255 | | 0.130 | 0.043 | DMN |
| No. 38 PostC_4 | | 0.244 | | 0.127 | 0.047 | DAN |
| No. 40 ParOper_1 | | 0.255 | | 0.133 | 0.047 | VAN |
| No. 186 pCunPCC_1 | | 0.200 | | 0.104 | 0.047 | DMN |
| ***Subcortical region*** | | | | | | |
| Hippocampus (R) | | 0.121 | | 0.104 | 0.253 | SUB |
| Amygdala (R) | | 0.028 | | 0.093 | 0.797 | SUB |
| Posterior Thalamus (R) | | -0.057 | | 0.103 | 0.621 | SUB |
| Anterior Thalamus (R) | | -0.044 | | 0.092 | 0.671 | SUB |
| Nucleus accumbens (R) | | -0.118 | | 0.082 | 0.147 | SUB |
| Globus pallidus (R) | | 0.021 | | 0.103 | 0.878 | SUB |
| Putamen (R) | | -0.088 | | 0.095 | 0.364 | SUB |
| Caudate nucleus (R) | | -0.083 | | 0.100 | 0.418 | SUB |
| Hippocampus (L) | | 0.036 | | 0.103 | 0.762 | SUB |
| Amygdala (L) | | 0.096 | | 0.087 | 0.280 | SUB |
| Posterior Thalamus (L) | | -0.089 | | 0.103 | 0.399 | SUB |
| Anterior Thalamus (L) | | -0.038 | | 0.093 | 0.719 | SUB |
| Nucleus accumbens (L) | | -0.114 | | 0.077 | 0.136 | SUB |
| Globus pallidus (L) | | 0.006 | | 0.101 | 0.958 | SUB |
| Putamen (L) | | -0.094 | | 0.090 | 0.308 | SUB |
| Caudate nucleus (L) | | -0.094 | | 0.096 | 0.337 | SUB |
| **Contrast: ISC_{low, high}_ > ISC_{low, low}_** | | | | | | |
| ***Significant cortical region*** | | | | | | |
| No. 138 PostC_2 | | 0.341 | | 0.147 | 0.020 | DAN |
| No. 137 PostC_1 | | 0.312 | | 0.140 | 0.024 | DAN |
| No. 2 ExStr_2 | | 0.280 | | 0.131 | 0.029 | VN |
| No. 29 TempOcc_1 | | 0.250 | | 0.121 | 0.035 | DAN |
| No.117 SomMotA_5 | | 0.284 | | 0.139 | 0.035 | SMN |
| No. 140 PostC_4 | | 0.276 | | 0.135 | 0.035 | DAN |
| No. 184 IPL_1 | | 0.224 | | 0.109 | 0.035 | DMN |
| No. 63 PFCl_1 | | -0.163 | | 0.082 | 0.035 | CN |
| No. 142 ParOper_1 | | 0.267 | | 0.138 | 0.043 | VAN |
| No. 135 SPL_3 | | 0.232 | | 0.123 | 0.049 | DAN |
| ***Subcortical region*** | | | | | | |
| Hippocampus (R) | | 0.114 | | 0.104 | 0.276 | SUB |
| Amygdala (R) | | 0.032 | | 0.093 | 0.762 | SUB |
| Posterior Thalamus (R) | | -0.072 | | 0.103 | 0.507 | SUB |
| Anterior Thalamus (R) | | -0.022 | | 0.092 | 0.844 | SUB |
| Nucleus accumbens (R) | | 0.097 | | 0.082 | 0.231 | SUB |
| Globus pallidus (R) | | 0.082 | | 0.103 | 0.427 | SUB |
| Putamen (R) | | -0.111 | | 0.095 | 0.245 | SUB |
| Caudate nucleus (R) | | -0.004 | | 0.100 | 0.969 | SUB |
| Hippocampus (L) | | 0.118 | | 0.103 | 0.252 | SUB |
| Amygdala (L) | | 0.105 | | 0.087 | 0.229 | SUB |
| Posterior Thalamus (L) | | 0.013 | | 0.103 | 0.913 | SUB |
| Anterior Thalamus (L) | | 0.008 | | 0.093 | 0.942 | SUB |
| Nucleus accumbens (L) | | -0.080 | | 0.077 | 0.300 | SUB |
| Globus pallidus (L) | | -0.056 | | 0.101 | 0.611 | SUB |
| Putamen (L) | | -0.066 | | 0.090 | 0.469 | SUB |
| Caudate nucleus (L) | | -0.069 | | 0.096 | 0.483 | SUB |

The index of cortical regions is taken from Schafer 200 atlas. FDR correction was applied to correct for multiple comparisons. All the reported *p*-values are two-tailed. R: right; L: left; DAN: dorsal attention network; CN: control network; DMN: default mode network; SMN: somatomotor network; SUB: subcortical network; VN: visual network. VAN: ventral attention network.

**Table S5. Dyad-level results that relate resilience similarity with ISC during neutral movie.**

| Region | β | SE | | Corrected  *p*-value | Network |
| --- | --- | --- | --- | --- | --- |
| ***Significant cortical regions*** |  | |  |  |  |
| No. 34 SPL_3 | 0.179 | | 0.040 | 0.002 | DAN |
| No. 141 FEF_1 | 0.171 | | 0.039 | 0.002 | DAN |
| No. 38 PostC_4 | 0.153 | | 0.039 | 0.006 | DAN |
| No. 40 ParOper_1 | 0.153 | | 0.038 | 0.006 | VAN |
| No. 43 FrOper_2 | 0.104 | | 0.027 | 0.008 | VAN |
| No. 137 PostC_1 | 0.149 | | 0.039 | 0.008 | DAN |
| No. 135 SPL_3 | 0.146 | | 0.039 | 0.008 | DAN |
| No. 127 S2_2 | 0.067 | | 0.019 | 0.016 | SMN |
| No. 28 Cent_2 | 0.088 | | 0.025 | 0.016 | SMN |
| No. 36 PostC_2 | 0.134 | | 0.040 | 0.016 | DAN |
| No. 134 SPL_2 | 0.134 | | 0.039 | 0.016 | DAN |
| No. 59 IPS_2 | 0.115 | | 0.035 | 0.022 | CN |
| No. 138 PostC_2 | 0.125 | | 0.039 | 0.022 | DAN |
| No. 14 SomMotA_2 | 0.064 | | 0.020 | 0.023 | SMN |
| No. 35 PostC_1 | 0.124 | | 0.040 | 0.023 | DAN |
| No. 45 FrMed_1 | -0.062 | | 0.019 | 0.023 | VAN |
| No. 60 IPS_3 | 0.124 | | 0.039 | 0.023 | CN |
| No. 107 ExStrInf_1 | 0.121 | | 0.038 | 0.023 | VN |
| No. 143 PrC_1 | 0.112 | | 0.034 | 0.023 | VAN |
| No. 140 PostC_4 | 0.123 | | 0.040 | 0.024 | DAN |
| No. 166 IPS_2 | 0.115 | | 0.039 | 0.032 | CN |
| No. 32 SPL_1 | 0.111 | | 0.039 | 0.047 | DAN |
| No. 37 PostC_3 | 0.112 | | 0.040 | 0.047 | DAN |
| ***Subcortical regions*** |  | |  |  |  |
| Hippocampus (R) | 0.038 | | 0.026 | 0.454 | SUB |
| Amygdala (R) | -0.009 | | 0.025 | 0.975 | SUB |
| Posterior Thalamus (R) | -0.029 | | 0.029 | 0.727 | SUB |
| Anterior Thalamus (R) | -0.035 | | 0.029 | 0.621 | SUB |
| Nucleus accumbens (R) | -0.066 | | 0.024 | 0.059 | SUB |
| Globus pallidus (R) | 0.017 | | 0.016 | 0.706 | SUB |
| Putamen (R) | 0.006 | | 0.019 | 0.975 | SUB |
| Caudate nucleus (R) | 0.008 | | 0.018 | 0.970 | SUB |
| Hippocampus (L) | 0.026 | | 0.030 | 0.791 | SUB |
| Amygdala (L) | -0.007 | | 0.024 | 0.975 | SUB |
| Posterior Thalamus (L) | -0.030 | | 0.029 | 0.706 | SUB |
| Anterior Thalamus (L) | -0.020 | | 0.026 | 0.842 | SUB |
| Nucleus accumbens (L) | -0.072 | | 0.026 | 0.059 | SUB |
| Globus pallidus (L) | 0.005 | | 0.016 | 0.975 | SUB |
| Putamen (L) | 0.025 | | 0.017 | 0.454 | SUB |
| Caudate nucleus (L) | -0.011 | | 0.019 | 0.909 | SUB |

The index of cortical regions is taken from Schafer 200 atlas. FDR correction was applied to correct for multiple comparisons. All the reported *p*-values are two-tailed. R: right; L: left; DAN: dorsal attention network; VAN: ventral attention network; CN: control network; VN: visual network; SMN: somatomotor network; SUB: subcortical network; VAN: ventral attention network..

**Table S6. Dyad-level results that related resilience similarity with ISC during negative movie.**

| Region | β | SE | | Corrected  *p*-value | Network |
| --- | --- | --- | --- | --- | --- |
| ***Significant cortical region*** |  | |  |  |  |
| No. 137 PostC_1 | 0.315 | | 0.036 | <0.001 | DAN |
| No. 138 PostC_2 | 0.283 | | 0.034 | <0.001 | DAN |
| No. 2 ExStr_2 | 0.260 | | 0.038 | <0.001 | VN |
| No. 104 ExStr_3 | 0.263 | | 0.039 | <0.001 | VN |
| No. 135 SPL_3 | 0.261 | | 0.039 | <0.001 | DAN |
| No. 26 S2_3 | 0.249 | | 0.039 | <0.001 | SMN |
| No. 129 S2_4 | 0.200 | | 0.033 | <0.001 | SMN |
| No. 5 ExStr_4 | 0.209 | | 0.039 | <0.001 | VN |
| No. 117 SomMotA_5 | 0.193 | | 0.037 | <0.001 | SMN |
| No. 113 SomMotA_1 | 0.180 | | 0.035 | <0.001 | SMN |
| No. 143 PrC_1 | 0.185 | | 0.037 | <0.001 | VAN |
| No. 64 PFCl_2 | 0.191 | | 0.039 | <0.001 | CN |
| No. 166 IPS_2 | 0.185 | | 0.039 | <0.001 | CN |
| No. 36 PostC_2 | 0.176 | | 0.038 | <0.001 | DAN |
| No. 141 FEF_1 | 0.180 | | 0.039 | <0.001 | DAN |
| No. 38 PostC_4 | 0.179 | | 0.040 | <0.001 | DAN |
| No. 24 S2_2 | 0.147 | | 0.032 | <0.001 | SMN |
| No. 37 PostC_3 | 0.155 | | 0.036 | <0.001 | DAN |
| No. 133 SPL_1 | 0.161 | | 0.038 | <0.001 | DAN |
| No. 146 FrOper_1 | 0.147 | | 0.033 | <0.001 | VAN |
| No. 39 FEF_1 | 0.150 | | 0.035 | <0.001 | DAN |
| No. 67 Temp_1 | 0.148 | | 0.035 | <0.001 | CN |
| No. 35 PostC_1 | 0.162 | | 0.039 | <0.001 | DAN |
| No. 132 ParOcc_1 | 0.157 | | 0.040 | 0.001 | DAN |
| No. 140 PostC_4 | 0.151 | | 0.038 | 0.001 | DAN |
| No. 18 SomMotA_6 | 0.153 | | 0.039 | 0.001 | SMN |
| No. 177 PFClv_1 | 0.146 | | 0.037 | 0.001 | CN |
| No. 147 FrMed_1 | 0.129 | | 0.033 | 0.001 | VAN |
| No. 112 ExStrSup_3 | 0.154 | | 0.040 | 0.001 | VN |
| No. 43 FrOper_2 | 0.129 | | 0.033 | 0.001 | VAN |
| No. 127 S2_2 | 0.152 | | 0.039 | 0.001 | SMN |
| No. 168 PFCl_1 | 0.147 | | 0.038 | 0.001 | CN |
| No. 86 Temp_4 | 0.147 | | 0.039 | 0.001 | DMN |
| No. 42 FrOper_1 | 0.139 | | 0.037 | 0.001 | VAN |
| No. 136 SPL_4 | 0.145 | | 0.039 | 0.002 | DAN |
| No. 70 PFClv_1 | 0.122 | | 0.033 | 0.002 | CN |
| No. 12 ExStrSup_2 | 0.147 | | 0.040 | 0.002 | VN |
| No. 68 IPL_1 | 0.130 | | 0.036 | 0.002 | CN |
| No. 105 ExStr_4 | 0.131 | | 0.037 | 0.002 | VN |
| No. 40 ParOper_1 | 0.138 | | 0.039 | 0.003 | VAN |
| No. 165 IPS_1 | 0.127 | | 0.036 | 0.003 | CN |
| No. 178 PFClv_2 | 0.131 | | 0.038 | 0.004 | CN |
| No. 52 OFC_2 | 0.126 | | 0.036 | 0.004 | LN |
| No. 115 SomMotA_3 | 0.118 | | 0.034 | 0.004 | SMN |
| No. 111 ExStrSup_2 | 0.136 | | 0.040 | 0.004 | VN |
| No. 32 SPL_1 | 0.131 | | 0.040 | 0.005 | DAN |
| No. 87 IPL_1 | 0.124 | | 0.038 | 0.007 | DMN |
| No. 182 pCun_2 | 0.125 | | 0.039 | 0.007 | CN |
| No. 79 pCunPCC_3 | 0.121 | | 0.038 | 0.007 | DMN |
| No. 157 OFC_1 | 0.113 | | 0.036 | 0.008 | LN |
| No. 62 PFClv_1 | 0.080 | | 0.026 | 0.012 | CN |
| No. 181 pCun_1 | 0.117 | | 0.040 | 0.014 | CN |
| No. 31 ParOcc_1 | 0.105 | | 0.036 | 0.014 | DAN |
| No. 16 SomMotA_4 | 0.094 | | 0.031 | 0.015 | SMN |
| No. 120 SomMotA_8 | 0.115 | | 0.040 | 0.015 | SMN |
| No. 109 StriCal_1 | 0.112 | | 0.039 | 0.019 | VN |
| No. 57 Temp_1 | 0.112 | | 0.040 | 0.019 | CN |
| No. 74 Cingp_1 | 0.108 | | 0.038 | 0.020 | CN |
| No. 75 IPL_1 | 0.106 | | 0.038 | 0.020 | DMN |
| No. 152 PFClv_1 | 0.088 | | 0.031 | 0.021 | VAN |
| No. 25 Aud_3 | 0.108 | | 0.040 | 0.023 | SMN |
| No. 175 PFCld_1 | 0.084 | | 0.031 | 0.027 | CN |
| No. 150 FrMed_2 | 0.094 | | 0.035 | 0.027 | VAN |
| No. 60 IPS_3 | 0.107 | | 0.040 | 0.027 | CN |
| No. 107 ExStrInf_1 | 0.105 | | 0.040 | 0.029 | VN |
| No. 158 OFC_2 | 0.099 | | 0.037 | 0.029 | LN |
| No. 119 SomMotA_7 | 0.075 | | 0.028 | 0.033 | SMN |
| No. 142 ParOper_1 | 0.098 | | 0.038 | 0.034 | VAN |
| No. 59 IPS_2 | 0.089 | | 0.034 | 0.034 | CN |
| No. 69 PFCl_1 | 0.081 | | 0.032 | 0.038 | CN |
| No. 176 PFCld_2 | 0.087 | | 0.035 | 0.045 | CN |
| No. 95 PFCv_4 | 0.091 | | 0.037 | 0.045 | DMN |
| ***Subcortical region*** |  | |  |  |  |
| Hippocampus (R) | 0.016 | | 0.038 | 0.816 | SUB |
| Amygdala (R) | 0.002 | | 0.035 | 0.978 | SUB |
| Posterior Thalamus (R) | 0.018 | | 0.038 | 0.804 | SUB |
| Anterior Thalamus (R) | 0.004 | | 0.035 | 0.971 | SUB |
| Nucleus accumbens (R) | 0.000 | | 0.031 | 0.997 | SUB |
| Globus pallidus (R) | 0.016 | | 0.038 | 0.816 | SUB |
| Putamen (R) | -0.015 | | 0.036 | 0.816 | SUB |
| Caudate nucleus (R) | -0.026 | | 0.037 | 0.663 | SUB |
| Hippocampus (L) | -0.027 | | 0.038 | 0.663 | SUB |
| Amygdala (L) | 0.044 | | 0.033 | 0.338 | SUB |
| Posterior Thalamus (L) | -0.018 | | 0.038 | 0.804 | SUB |
| Anterior Thalamus (L) | -0.030 | | 0.035 | 0.583 | SUB |
| Nucleus accumbens (L) | -0.036 | | 0.029 | 0.385 | SUB |
| Globus pallidus (L) | -0.004 | | 0.037 | 0.971 | SUB |
| Putamen (L) | -0.019 | | 0.034 | 0.758 | SUB |
| Caudate nucleus (L) | 0.007 | | 0.036 | 0.923 | SUB |

The index of cortical regions is taken from Schafer 200 atlas. FDR correction was applied to correct for multiple comparisons. All the reported *p*-values are two-tailed. R: right; L: left; DAN: dorsal attention network; CN: control network; DMN: default mode network; SMN: somatomotor network; SUB: subcortical network; VN: visual network. LN: limbic network; VAN: ventral attention network.

**Table S7. The modulation effects of intolerance of uncertainty on resilience-driven neural synchrony in the neutral movie.**

| Region | β | SE | | Corrected  *p*-value | Network |
| --- | --- | --- | --- | --- | --- |
| ***Significant cortical region*** |  | |  |  |  |
| No. 28 Cent_2 | -0.096 | | 0.018 | 0.000 | SMN |
| No. 143 FrOper_2 | -0.082 | | 0.018 | 0.001 | VAN |
| No. 37 PostC_3 | -0.064 | | 0.015 | 0.002 | DAN |
| No. 140 PostC_4 | -0.067 | | 0.016 | 0.002 | DAN |
| No. 150 FrMed_2 | -0.074 | | 0.018 | 0.002 | VAN |
| No. 134 SPL_2 | -0.069 | | 0.017 | 0.002 | DAN |
| No. 114 SomMotA_2 | -0.070 | | 0.018 | 0.003 | DAN |
| No. 198 TempPar_2 | -0.067 | | 0.018 | 0.006 | TempPar |
| No. 123 SomMotA_11 | -0.066 | | 0.018 | 0.006 | SMN |
| No. 99 TempPar_1 | -0.066 | | 0.018 | 0.006 | TempPar |
| No. 20 SomMotA_8 | -0.063 | | 0.018 | 0.009 | SMN |
| No. 22 Aud_2 | -0.062 | | 0.018 | 0.009 | SMN |
| No. 12 ExStrSup_2 | -0.056 | | 0.016 | 0.009 | VN |
| No. 139 PostC_3 | -0.060 | | 0.018 | 0.009 | DAN |
| No. 38 PostC_4 | -0.055 | | 0.017 | 0.013 | DAN |
| No. 15 SomMotA_3 | -0.059 | | 0.018 | 0.015 | SMN |
| No. 19 SomMotA_7 | -0.057 | | 0.018 | 0.019 | SMN |
| No. 136 SPL_4 | -0.051 | | 0.016 | 0.020 | DAN |
| No. 119 SomMotA_7 | -0.056 | | 0.018 | 0.024 | SMN |
| No. 27 Cent_1 | -0.054 | | 0.018 | 0.024 | SMN |
| No. 166 IPS_2 | -0.052 | | 0.017 | 0.024 | CN |
| No. 40 ParOper_1 | -0.051 | | 0.017 | 0.029 | DAN |
| No. 44 ParMed_1 | -0.054 | | 0.018 | 0.029 | VAN |
| No. 35 PostC_1 | -0.047 | | 0.016 | 0.033 | VAN |
| No. 17 SomMotA_5 | -0.052 | | 0.018 | 0.033 | SMN |
| No. 137 PostC_1 | -0.043 | | 0.015 | 0.033 | DAN |
| No. 199 TempPar_3 | -0.053 | | 0.018 | 0.033 | TempPar |
| No. 49 Ins_1 | -0.049 | | 0.017 | 0.036 | VAN |
| No. 141 FEF_1 | -0.050 | | 0.018 | 0.040 | DAN |
| ***Subcortical region*** |  | |  |  |  |
| Hippocampus (R) | 0.038 | | 0.026 | 0.454 | SUB |
| Amygdala (R) | -0.009 | | 0.025 | 0.975 | SUB |
| Posterior Thalamus (R) | -0.029 | | 0.029 | 0.727 | SUB |
| Anterior Thalamus (R) | -0.035 | | 0.029 | 0.621 | SUB |
| Nucleus accumbens (R) | -0.066 | | 0.024 | 0.059 | SUB |
| Globus pallidus (R) | 0.017 | | 0.016 | 0.706 | SUB |
| Putamen (R) | 0.006 | | 0.019 | 0.975 | SUB |
| Caudate nucleus (R) | 0.008 | | 0.018 | 0.970 | SUB |
| Hippocampus (L) | 0.026 | | 0.030 | 0.791 | SUB |
| Amygdala (L) | -0.007 | | 0.024 | 0.975 | SUB |
| Posterior Thalamus (L) | -0.030 | | 0.029 | 0.706 | SUB |
| Anterior Thalamus (L) | -0.020 | | 0.026 | 0.842 | SUB |
| Nucleus accumbens (L) | -0.072 | | 0.026 | 0.059 | SUB |
| Globus pallidus (L) | 0.005 | | 0.016 | 0.975 | SUB |
| Putamen (L) | 0.025 | | 0.017 | 0.454 | SUB |
| Caudate nucleus (L) | -0.011 | | 0.019 | 0.909 | SUB |

The index of cortical regions is taken from Schafer 200 atlas. FDR correction was applied to correct for multiple comparisons. All the reported *p*-values are two-tailed. R: right; L: left; DAN: dorsal attention network; CN: control network; VN: visual network; SMN: somatomotor network; SUB: subcortical network; VAN: ventral attention network; TempPar: temporal parietal area.

**Table S8. The modulation effects of intolerance of uncertainty on resilience-driven neural synchrony in the negative movie.**

| Region | β | SE | | Corrected  *p*-value | Network |
| --- | --- | --- | --- | --- | --- |
| ***Significant cortical region*** |  | |  |  |  |
| No. 14 SomMotA_2 | -0.092 | | 0.018 | <0.001 | SMN |
| No. 35 PostC_1 | -0.080 | | 0.016 | <0.001 | DAN |
| No. 38 PostC_4 | -0.079 | | 0.016 | <0.001 | DAN |
| No. 36 PostC_2 | -0.058 | | 0.014 | 0.003 | DAN |
| No. 139 PostC_3 | -0.067 | | 0.017 | 0.004 | DAN |
| No. 74 Cingp_1 | 0.066 | | 0.017 | 0.004 | CN |
| No. 5 ExStr_4 | -0.055 | | 0.014 | 0.005 | VN |
| No. 126 S2_1 | 0.066 | | 0.018 | 0.006 | SMN |
| No. 156 PFCmp_1 | 0.062 | | 0.017 | 0.009 | VAN |
| No. 50 PFCmp_1 | 0.060 | | 0.017 | 0.012 | VAN |
| No. 61 PFCd_1 | -0.061 | | 0.018 | 0.014 | CN |
| No. 33 SPL_2 | -0.057 | | 0.017 | 0.014 | DAN |
| No. 37 PostC_3 | -0.042 | | 0.013 | 0.014 | DAN |
| No. 82 PFCm_3 | 0.059 | | 0.018 | 0.014 | DMN |
| No. 78 pCunPCC_2 | 0.057 | | 0.017 | 0.015 | DMN |
| No. 72 pCun_1 | 0.055 | | 0.017 | 0.016 | CN |
| No. 167 PFCd_1 | -0.058 | | 0.018 | 0.017 | CN |
| No. 26 S2_3 | -0.050 | | 0.016 | 0.022 | SMN |
| No. 92 PFCv_1 | 0.054 | | 0.017 | 0.022 | DMN |
| No. 166 IPS_2 | -0.051 | | 0.016 | 0.024 | CN |
| No. 67 Temp_1 | 0.054 | | 0.018 | 0.024 | CN |
| No. 88 PFCd_1 | 0.054 | | 0.018 | 0.024 | DMN |
| No. 181 pCun_1 | 0.049 | | 0.016 | 0.025 | CN |
| No. 154 Ins_1 | 0.049 | | 0.016 | 0.026 | VAN |
| No. 140 PostC_4 | -0.041 | | 0.014 | 0.033 | DAN |
| No. 68 IPL_1 | 0.049 | | 0.018 | 0.041 | CN |
| ***Subcortical region*** |  | |  |  |  |
| Hippocampus (R) | 0.016 | | 0.017 | 0.622 | SUB |
| Amygdala (R) | 0.001 | | 0.018 | 0.995 | SUB |
| Posterior Thalamus (R) | -0.017 | | 0.018 | 0.608 | SUB |
| Anterior Thalamus (R) | -0.013 | | 0.018 | 0.696 | SUB |
| Nucleus accumbens (R) | 0.052 | | 0.018 | 0.036 | SUB |
| Globus pallidus (R) | 0.041 | | 0.018 | 0.102 | SUB |
| Putamen (R) | 0.007 | | 0.018 | 0.836 | SUB |
| Caudate nucleus (R) | 0.016 | | 0.018 | 0.622 | SUB |
| Hippocampus (L) | 0.031 | | 0.018 | 0.246 | SUB |
| Amygdala (L) | 0.031 | | 0.018 | 0.264 | SUB |
| Posterior Thalamus (L) | 0.011 | | 0.018 | 0.731 | SUB |
| Anterior Thalamus (L) | 0.008 | | 0.018 | 0.826 | SUB |
| Nucleus accumbens (L) | 0.029 | | 0.018 | 0.302 | SUB |
| Globus pallidus (L) | 0.021 | | 0.018 | 0.528 | SUB |
| Putamen (L) | 0.006 | | 0.018 | 0.855 | SUB |
| Caudate nucleus (L) | 0.011 | | 0.018 | 0.734 | SUB |

The index of cortical regions is taken from Schafer 200 atlas. FDR correction was applied to correct for multiple comparisons. All the reported *p*-values are two-tailed. R: right; L: left; DAN: dorsal attention network; CN: control network; DMN: default mode network; SMN: somatomotor network; SUB: subcortical network; VN: visual network; VAN: ventral attention network.
